# RNA assignment and segmentation geometry in Visium HD profiles of fibrostenotic Crohn’s disease

**DOI:** 10.64898/2026.10.27.762235

**Authors:** Luca Massimino, Federica Ungaro

## Abstract

Retrospective reuse of spatial transcriptomics often starts from a deposited bin-level Seurat object rather than the inputs needed to revisit upstream molecular processing. We tested whether image segmentation could support a defensible cell-like reconstruction under this constraint in two complete H&E-stained intestinal sections from a Visium HD study of Crohn’s disease-associated fibrostenosis (GSE302569). The available data comprised 18,085 genes, 988,942 original 8 µm bins and 179,596,575 UMIs. We evaluated 20 segmentation configurations in the preserved image coordinate system using a common bin-centre assignment rule, without restarting Space Ranger or introducing higher-resolution expression. RNA assignment ranged from 0% to 74.68% in CD1 and 0% to 64.05% in CD2. Expansion increased recovery but did not eliminate merged regions, straight boundaries or within-bin ambiguity; approximately 70-73% of InstanSeg brightfield nuclear regions received no assigned RNA. All 38 non-empty configuration-slice combinations underwent native full-mode RCTD against the same nine-group scIBD reference. Mean slice-level RNA recovery was negatively associated with primary-component dominance across 19 configurations (descriptive Spearman ρ = −0.77). Neither dominant-call agreement nor concentrated RCTD support validates boundaries or cell identity. The observed limitations in RNA recovery, region geometry, and compositional support did not establish any tested configuration as a reliable single-cell reconstruction from these inputs. The findings identify limitations of retrospective single-cell reconstruction from the available binned inputs, not the impossibility of cell-resolved analysis from suitable original data.

## Introduction

Spatial transcriptomics preserves tissue context while measuring RNA across a section. Image-derived polygons can provide a useful unit for aggregating expression, but their biological interpretation depends on what was segmented and the spatial resolution of the available RNA. A nuclear region, an expanded nuclear territory, and a proposed whole cell are different targets. Assigning more RNA to one of these targets does not by itself establish that its boundaries are more accurate.

The expanding availability of spatial transcriptomic datasets creates opportunities to revisit existing studies at finer anatomical resolution. However, datasets originally analysed using spot- or bin-based frameworks do not necessarily include validated cell boundaries or cell-resolved expression matrices, limiting their direct reuse for single-cell-level spatial analyses. Computational segmentation and spatial aggregation methods seek to bridge this gap by associating deposited expression measurements with image-derived regions [3–7]. Their practical utility nevertheless depends on both the resolution of the available molecular data and the ability to delineate biologically meaningful structures. In architecturally heterogeneous tissues, a configuration suitable for one compartment may under-segment, over-segment or misrepresent another; retrospective conversion therefore requires tissue-aware evaluation rather than assuming that the availability of a segmentation tool is sufficient.

Intestinal tissue poses a particularly demanding setting for image-based partitioning. Tightly packed epithelial cells surround crypt and villus profiles, while immune and mesenchymal populations occupy the lamina propria and deeper layers contain vascular, stromal and smooth-muscle structures with different shapes and packing densities. Human intestinal atlases describe spatially organised cellular neighbourhoods rather than a morphologically uniform sheet [8,9]. A configuration that separates nuclear profiles in one compartment may therefore behave differently at epithelial-stromal interfaces or within elongated myoid fields.

The source study characterizes Crohn’s disease-associated fibrostenosis using high-resolution spatial transcriptomics [1,2]. The publicly available dataset permits evaluation of segmentation across intestinal tissue containing dense mucosal and immune-rich fields, stromal regions and muscular structures. The expression data analysed here are provided as 8 µm bins. Expression can therefore be aggregated by assigning complete bin-level count vectors to segmented regions, but individual transcript molecules cannot be localized within each bin.

Fibrostenotic Crohn’s disease adds architectural remodelling to this baseline heterogeneity. The source study describes submucosal expansion of extracellular-matrix-producing myofibroblasts and altered immune-stromal organisation [1]. Dense inflammatory fields, glandular profiles and collagen-rich or muscularised regions consequently provide different image contexts. These features motivate treating conversion to polygon-associated expression as a non-trivial, tissue-specific measurement problem, rather than assuming that one checkpoint or fixed expansion generates equivalent biological units throughout the section.

A recent spatial transcriptomics segmentation benchmark provides relevant methodological context [3]. Here, we address a narrower practical question: can selected image-segmentation configurations support a convincing retrospective conversion to cell-like expression units when only a deposited 8 µm bin-level Seurat object and its embedded H&E are used, without restarting from Space Ranger or finer-resolution molecular inputs? We evaluated Cellpose configurations, InstanSeg, microscopy and histopathology SAM configurations, and classical watershed variants [4–7]. We assessed RNA recovery, qualitative morphology, and downstream RCTD compositional support separately, retaining sparse, excluded and zero-region outcomes rather than combining them into an accuracy score. Figure 1 outlines this deposited-input workflow.

**Figure 1.**
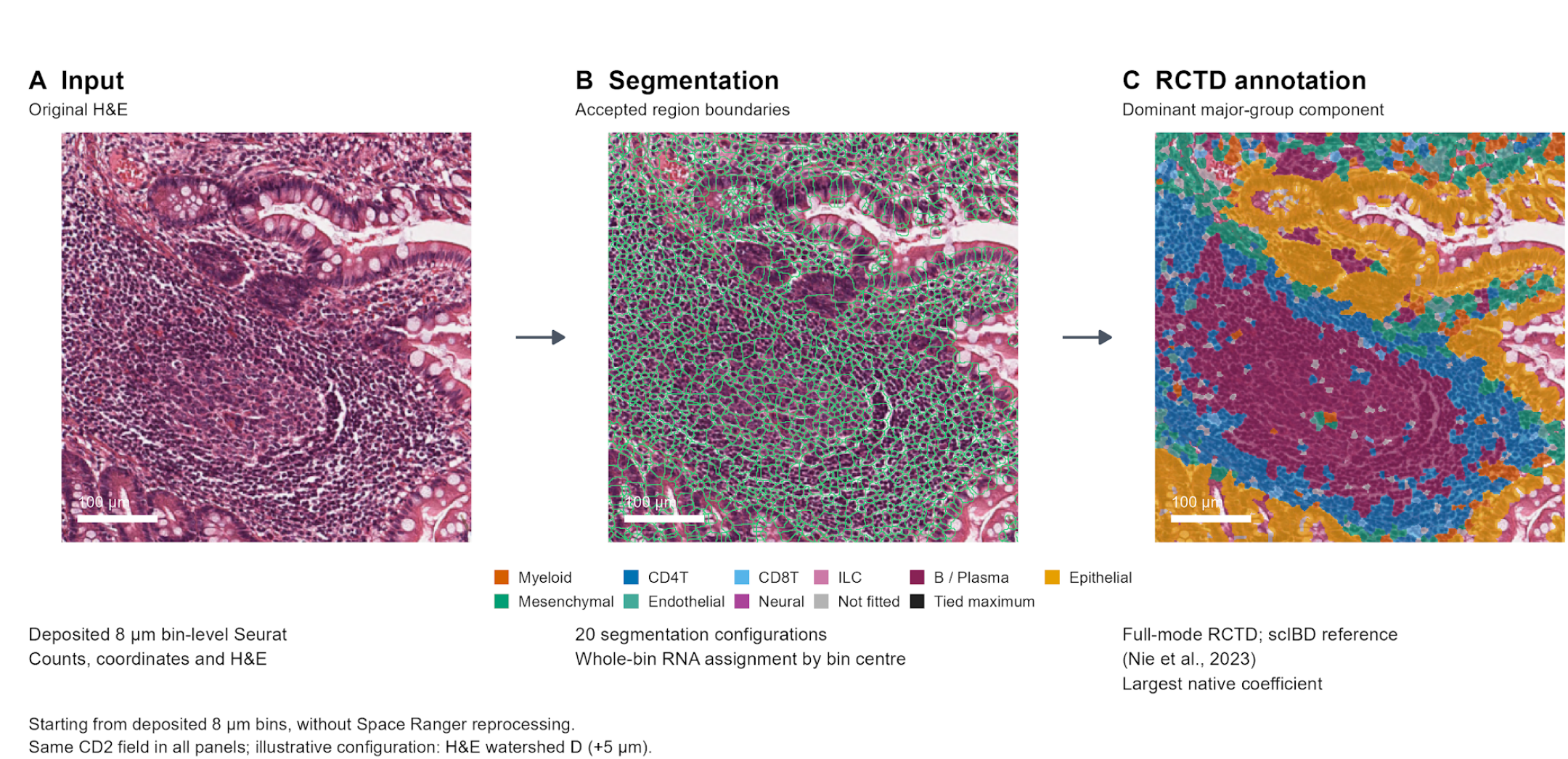
Workflow from deposited 8 µm bin-level Seurat to segmentation-derived regions and RCTD major-group display calls. The same CD2 field is shown throughout; no Space Ranger reprocessing or finer-resolution RNA information is introduced.

## Methods

### Data and coordinate preservation

We started from the deposited bin-level Seurat object for CD1 and CD2 (GSE302569), omprising 520,101 and 468,841 original 8 µm bins, respectively. Count values were not modified; CD1 contained 103,577,252 UMIs, and CD2 contained 76,019,323 UMIs. Analysis was deliberately restricted to these available counts, coordinates, and embedded H&E images: we did not reprocess raw sequencing through Space Ranger or substitute finer-resolution molecular measurements. The histological images stored in the VisiumV2 objects measured 6,000 by 4,792 and 6,000 by 5,148 pixels. Full-resolution bin coordinates were transformed to the deposited image coordinate system using the stored high-resolution image scale factor. No rotation, reflection, registration adjustment or coordinate refitting was applied. Physical pixel sizes estimated from the original 8 µm lattice were 1.4 and 1.3 µm per pixel; these values supported physical-size parameters and area diagnostics rather than replacing the original coordinates.

### Segmentation configurations

The ten learned configurations were Cellpose cyto3, Cellpose cpsam, cpsam_v2, cpdino-vitb and cpdino; InstanSeg fluorescence_nuclei_and_cells and brightfield_nuclei; microSAM vit_b_lm with automatic instance segmentation (AIS) or automatic mask generation (AMG); and PathoSAM vit_b_histopathology with AIS. These are specific checkpoint and interface configurations, not ten equivalent biological targets. The fluorescence InstanSeg model was retained explicitly as an out-of-domain comparison on H&E.

Cellpose cyto3 received a bright hematoxylin signal derived from the original H&E, with its 99th percentile within the common tissue mask mapped to 255 and clipped to uint8. The other Cellpose configurations received original RGB H&E. The diameter parameter was 20 pixels, and the batch size was one. No image-restoration model or dataset-specific Cellpose training was used. InstanSeg received float32 RGB input, the calibrated pixel size, native 512-pixel tiling, and batch size one, with the corresponding cell or nuclear prediction target and rescaling back to the original frame. SAM AIS used original RGB, 384-pixel tiles, and 64-pixel halos. The LM AMG configuration used grayscale H&E, 1200-pixel tiles, 128-pixel halos, predicted IoU threshold 0.88, stability threshold 0.95, and points per batch eight. The software-provided tiling and inference interfaces were used; learned output masks were not expanded or custom-thresholded after inference.

The classical configurations comprised scikit-image and OpenCV baselines and eight marker-controlled hematoxylin watershed configurations implemented with scikit-image. Configurations A–D crossed hole-filling rules and nominal seed separation; each was evaluated without expansion and with 5 micrometre expansion (Supplementary Table S1). Hematoxylin was separated from RGB H&E using rgb2hed, smoothed, and thresholded by Otsu within the common tissue mask. For A–D, markers were local maxima of the foreground Euclidean distance transform, and watershed was applied to the negative distance map within that foreground. Label expansion used expand_labels and was restricted to the same tissue mask. Exact preprocessing and parameter values are given in Supplementary Table S1. The two baseline implementations used Gaussian sigma one pixel, minimum foreground object size 15 pixels, and a peak separation of ten pixels. All exact settings and code revisions are archived.

### RNA assignment and outcome definitions

Each configuration was evaluated across both complete tissue sections. For each original bin, its scaled centre coordinates were floored to the image pixel grid. If that pixel carried a positive region label, the entire original gene-count vector was assigned to that region; otherwise, the bin remained unassigned. The same accepted mappings were used for RNA accounting and subsequent gene-by-region aggregation, without reallocating fractions of a bin between regions. Assigned and unassigned UMI totals were required to equal the original section total exactly. Outside-image centres remained unassigned: 397 bins and 655 UMIs in CD1, and 262 bins and 175 UMIs in CD2.

We reported assigned RNA and bins, the number of distinct positive region labels, regions with no assigned RNA, region areas, UMI and bin distributions per region, image and common tissue coverage, and execution-resource measurements. A common H&E-derived tissue mask was used only as a shared diagnostic denominator and where explicitly required by classical configurations. Bin-corner discordance measured whether any of the four original lattice-derived bin corners had a label different from the centre label. Its UMI-weighted form reports RNA in such bins; it is not an estimate of misassigned molecules. Zero-region outputs have undefined per-region distributions rather than artificial zero-valued distributions.

### Reproducibility and visual review

Versions included Cellpose 3.1.1.2 and 4.2.1, instanseg-torch 0.1.2, micro-sam 1.8.14 and Torch 2.5.1 with CUDA 12.1. Inference used fixed random seeds, recorded model hashes and native output readback. InstanSeg input was represented as float32 to satisfy the inference interface. Outputs from unsuccessful executions were excluded; only outputs from complete, successful reruns were analysed. Computational consistency checks verified all 40 accepted masks, source identity, bin-centre label assignments and exact UMI conservation. Timing was measured once per section and configuration, with model loading reported separately. Concurrent execution and different inference interfaces preclude interpreting these timings as a controlled speed benchmark.

Comparison panels show each entire section and the same dense and medium-density fields selected from H&E independently of segmentation. All detected boundaries are shown, with Cellpose configurations first. Figure 1 illustrates the workflow in the preselected dense CD2 field using H&E watershed D (+5 µm), the same example shown in Supplementary Figures S8 and S16; it is an illustration, not a selected best-performing configuration. Visual review was qualitative and unblinded, without manually annotated ground truth, inter-rater assessment or quantitative boundary accuracy. RCTD was subsequently used as a downstream compositional assessment of the accepted count aggregations, separately from this visual boundary review.

### Polygon-associated expression and RCTD analysis

For downstream compositional analysis, complete gene-count vectors from the original 8 µm bins were summed using the accepted bin-to-region mappings. All detected regions, including zero-RNA regions, were retained in the region accounting. Assigned and unassigned expression was checked against the original counts, including exact gene-wise conservation. This aggregation changes the unit of analysis but does not recover the position or cellular origin of molecules within a bin.

The common reference was the Crohn’s ileum scIBD reference of Nie et al. [10], with original integer RNAraw counts and author-defined major_cluster labels: Myeloid, CD4T, CD8T, ILC, B_Plasma, Epithelial, Mesenchymal, Endothelial and Neural. Each non-empty configuration–slice combination was analysed with the native Bioconductor spacexr 1.4.0 createRctd/runRctd interface in full mode with native filtering [11,12]. Native fitted objects, coefficients, metadata and exclusions were preserved. No fit was fabricated for zero-region outputs, and filtered regions were not removed from segmentation or RNA-recovery accounting. Full mode does not restrict each region to one or two cell types.

### Concordance, ranked contributions and recovery–dominance comparison

For the dominant-major-group maps, the group with the largest native full-mode coefficient defines a display call; these calls are not calibrated posterior probabilities. For pairwise concordance, each fitted region’s call was mapped back through the original bin-to-region assignment. Within each slice, agreement was the fraction of original bins with identical dominant labels among bins fitted in both configurations. Bins absent or filtered in either method were excluded from that pair’s denominator rather than counted as matching labels. The two slice-specific matrices were averaged arithmetically; average-linkage dendrograms used distance 1 minus mean agreement.

For the ranked-contribution display, each fitted polygon’s nine native coefficients were divided by their unmodified sum and sorted from largest (Call 1) to smallest (Call 9), independently of identity. Ranked contributions were averaged across fitted polygons within each slice and then averaged arithmetically across slices. This display-only scaling did not clip, threshold or overwrite native coefficients. Call 1–9 denote ordered contributions, not distinct statistically accepted identities; no identity-count threshold or formal uncertainty score was applied. For Figure 3C, the x coordinate was the arithmetic mean of the two assigned/original UMI fractions, using exactly the complete-section denominator of Figure 2; the y coordinate was the same mean Call 1 contribution as in Figure 3B. Neither raw UMIs nor polygons were pooled across slices for these averages. Spearman correlation was descriptive across the 19 configurations with non-empty fits, without an inferential P value.

**Figure 2.**
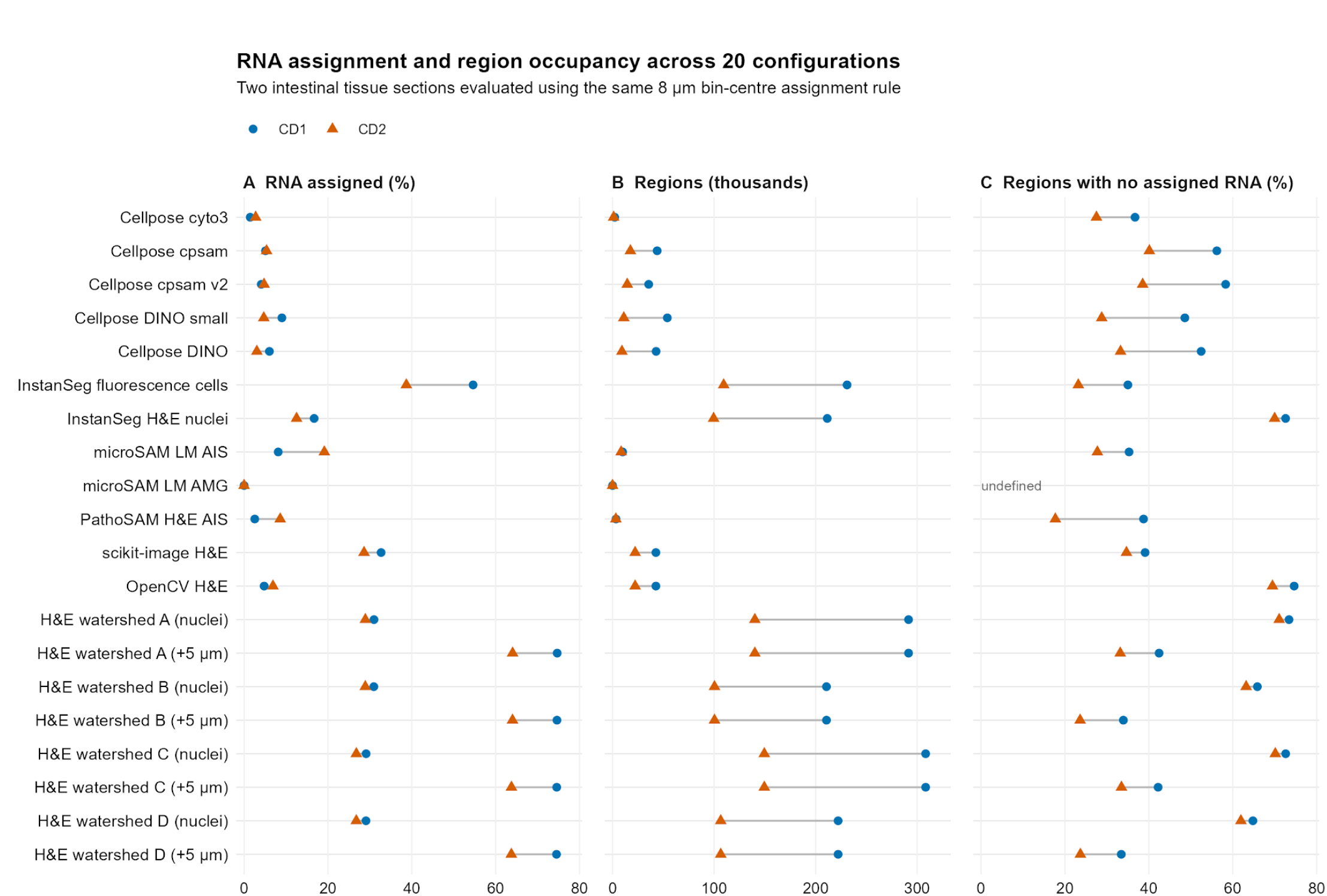
RNA assignment, detected region counts and RNA occupancy across CD1 and CD2. Greater RNA coverage does not establish more accurate cellular boundaries. Method configurations and complete diagnostic metrics are provided in Supplementary Table S1.

**Figure 3.**
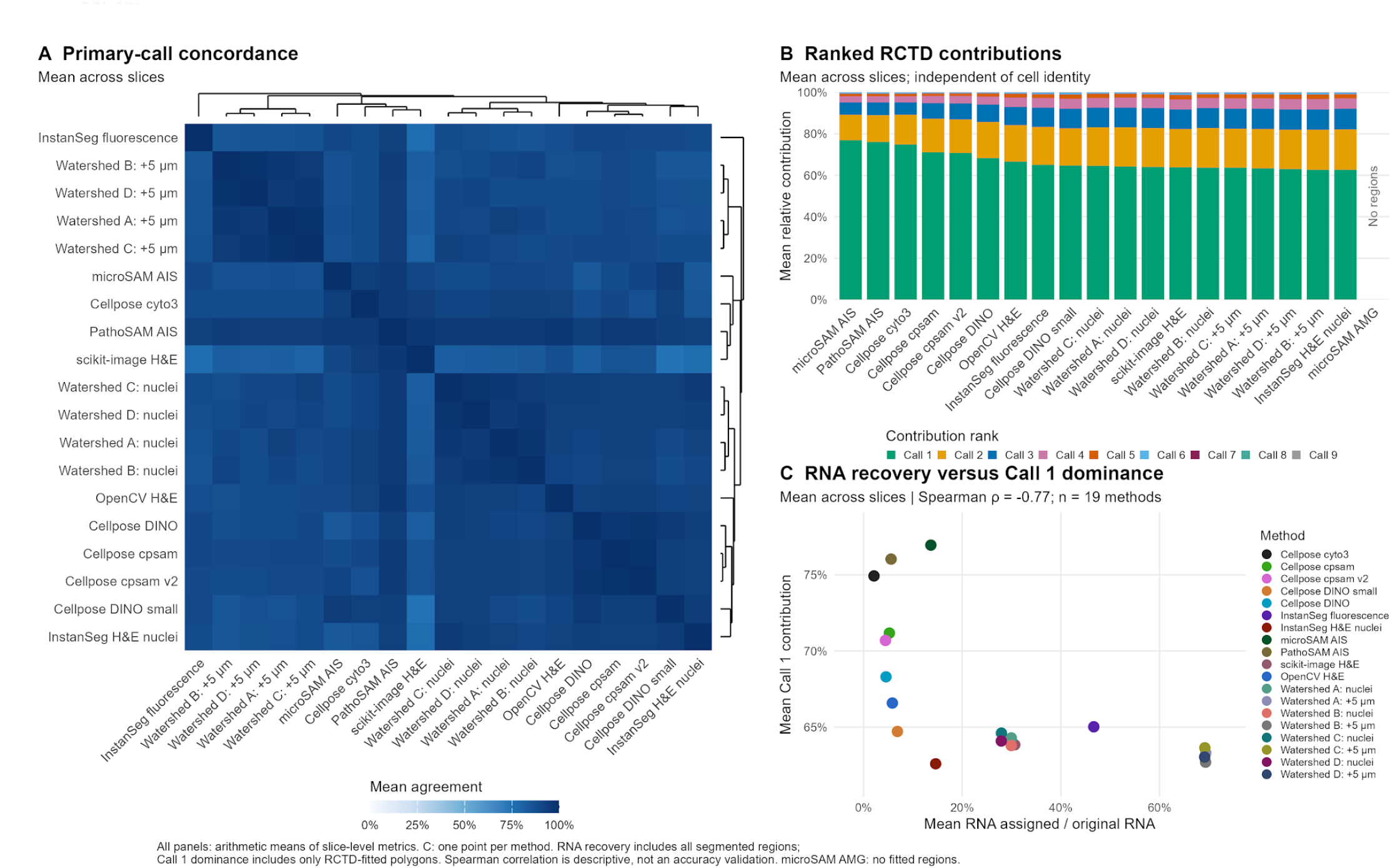
Concordance and ranked RCTD contributions across segmentation configurations. All summaries are arithmetic means of slice-level metrics. C shows the trade-off between RNA recovery and Call 1 dominance (Spearman ρ = −0.77; 19 configurations). See the full legend above for denominators and interpretation.

## Results

### RNA assignment varied strongly across configurations

RNA assignment varied markedly across the 20 segmentation configurations, ranging from 0% to 74.68% in CD1 and from 0% to 64.05% in CD2 (Figure 2; Table 1; Supplementary Table S1). Cellpose cyto3 assigned 1.45% of CD1 RNA and 2.76% of CD2 RNA. Other tested Cellpose configurations assigned 4.09-9.02% in CD1 and 3.04-5.37% in CD2 (Supplementary Figures S1 and S5). InstanSeg brightfield nuclei assigned 17,307,900 and 9,517,888 UMIs, equivalent to 16.71% and 12.52%. Its fluorescence cell-target configuration assigned 54.62% and 38.70%, but that difference combines a change in biological target with an out-of-domain model and cannot be read as greater accuracy (Figure 2A; Supplementary Figures S2 and S6).

**Table 1.** RNA assignment and region counts.

| Configuration | CD1 RNA % | CD2 RNA % | CD1 regions | CD2 regions |
|---|---|---|---|---|
| Cellpose cyto3 | 1.45 | 2.76 | 2,174 | 1,094 |
| Cellpose cpsam | 5.11 | 5.37 | 43,931 | 17,594 |
| Cellpose cpsam v2 | 4.09 | 4.76 | 35,598 | 14,487 |
| Cellpose DINO small | 9.02 | 4.70 | 53,964 | 11,104 |
| Cellpose DINO | 6.04 | 3.04 | 42,902 | 9,247 |
| InstanSeg fluorescence cells | 54.62 | 38.70 | 230,986 | 109,464 |
| InstanSeg H&E nuclei | 16.71 | 12.52 | 211,462 | 99,547 |
| microSAM LM AIS | 8.14 | 19.14 | 9,934 | 8,477 |
| microSAM LM AMG | 0.00 | 0.00 | 0 | 0 |
| PathoSAM H&E AIS | 2.55 | 8.61 | 3,418 | 3,246 |
| scikit-image H&E | 32.68 | 28.61 | 42,695 | 22,250 |
| OpenCV H&E | 4.78 | 6.89 | 42,695 | 22,250 |
| H&E watershed A (nuclei) | 30.98 | 28.92 | 291,533 | 140,149 |
| H&E watershed A (+5 µm) | 74.68 | 64.05 | 291,533 | 140,149 |
| H&E watershed B (nuclei) | 30.95 | 28.91 | 210,697 | 100,416 |
| H&E watershed B (+5 µm) | 74.63 | 64.03 | 210,697 | 100,416 |
| H&E watershed C (nuclei) | 29.10 | 26.78 | 308,393 | 149,465 |
| H&E watershed C (+5 µm) | 74.57 | 63.77 | 308,393 | 149,465 |
| H&E watershed D (nuclei) | 29.07 | 26.76 | 222,174 | 106,576 |
| H&E watershed D (+5 µm) | 74.52 | 63.75 | 222,174 | 106,576 |

Percentages use the complete original section UMI totals. Region counts are not independently validated cell counts. Complete diagnostics and method configurations are provided in Supplementary Table S1.

The microSAM LM AIS configuration assigned 8.14% and 19.14% of section UMIs to regions in CD1 and CD2, respectively; the corresponding values for PathoSAM H&E AIS were 2.55% and 8.61% (Figure 2A; Supplementary Figures S2 and S6). The tested microSAM LM AMG configuration completed with zero positive regions in both sections (Figure 2B). This null result is retained rather than replaced by another configuration. It describes the tested checkpoint, grayscale input and thresholds, not the general capability of SAM or microSAM.

### Expansion increased RNA assignment without validating boundaries

The scikit-image H&E baseline assigned 32.7% and 28.6%, whereas the OpenCV H&E baseline assigned 4.8% and 6.9%. The unexpanded H&E watershed configurations were assigned approximately 29-31% in CD1 and 27-29% in CD2. Five micrometre expansion increased those ranges to approximately 74.5-74.7% and 63.7-64.1% (Figure 2A; Supplementary Figures S3, S4, S7 and S8). H&E watershed A increased from 32,092,775 to 77,353,849 assigned UMIs in CD1 and from 21,987,652 to 48,690,362 in CD2, without changing its region count (Figure 2B; Table 1). Across the two sections, its expanded regions received 126,044,211 of 179,596,575 UMIs, or approximately 70.18%.

The visual panels also show failure modes that this RNA increase cannot exclude. The illustrative CD2 field in Figure 1A–C provides a direct view of the original H&E, accepted boundaries and downstream RCTD display calls. Several H&E watershed configurations contain large merged regions and long straight boundaries, particularly in the dense CD2 field (Supplementary Figures S7 and S8). The scikit-image H&E baseline occasionally merged adjacent nuclear structures (Supplementary Figures S3 and S7). Several learned configurations omit dense nuclear fields or outline pale luminal or goblet-associated structures (Supplementary Figures S1, S2, S5 and S6). These qualitative observations show why RNA assignment alone is insufficient to select a segmentation configuration for downstream polygon-based analysis (Supplementary Figures S1–S8).

### Many nuclear regions received no assigned RNA at the available bin resolution

The InstanSeg brightfield nuclear model produced 211,462 regions in CD1 and 99,547 in CD2, but 72.6% and 70.0% received zero UMIs under the common assignment rule (Figure 2B–C; Supplementary Table S1). This endpoint combines segmentation geometry, bin size, centre placement and expression sparsity. A region without assigned RNA is not necessarily a false nucleus. Conversely, assigning a complete 8 micrometre bin to a small region does not establish that all RNA in that bin originated from the corresponding nucleus or cell.

For expanded H&E watershed B, UMI-weighted bin-corner discordance was 83.7% and 71.2%. The fluorescence InstanSeg configuration had values of 88.2% and 73.6% (Supplementary Table S1). Thus, many RNA-bearing bins intersect label transitions under these geometries. These percentages are indicators of sensitivity to bin-centre assignment, not segmentation error rates or estimates of the number of incorrectly allocated molecules.

### Cross-method agreement and compositional dominance answer different questions

Full-mode RCTD was completed for all 38 non-empty configuration–slice combinations using the same nine-group scIBD reference. microSAM AMG had no regions in either slice and therefore no fit. Dominant-major-group maps are shown for Cellpose (Supplementary Figures S9 and S13), InstanSeg and the SAM-based configurations (Supplementary Figures S10 and S14), and the classical baselines and watershed configurations (Supplementary Figures S11, S12, S15 and S16). Regions excluded by the native filters remain visible as excluded rather than being assigned an artificial identity.

Figure 3A summarizes dominant-call concordance on a common molecular support: original bins assigned to fitted regions by both configurations in each comparison. Agreement on this conditional support does not imply that the methods recovered the same amount of tissue RNA or delineated the same cellular boundaries. Figure 3B separates the magnitude of the primary, secondary and lower-ranked contributions from which major group occupied each rank. A dominant Call 1 indicates concentrated compositional support; distributed ranks indicate a less concentrated fit, not a calibrated probability that a cell was misclassified. A multi-component fit alone cannot determine whether a region contains multiple cells, whether one cell has an ambiguous reference match, or whether aggregation and count depth have broadened its support.

### RNA recovery and primary-component dominance were inversely associated

Across the 19 non-empty configurations, mean slice-level RNA recovery was negatively associated with mean Call 1 contribution (descriptive Spearman ρ = −0.77; Figure 3C). Methods with high dominance could recover only a small fraction of the original RNA, while expanded watershed configurations recovered substantially more RNA but showed less concentrated RCTD support. The association does not establish a causal effect of recovery on dominance or create a single winner: the two axes have different inclusion denominators, the configurations are not independent biological replicates, and neither axis measures ground-truth cellular accuracy.

## Discussion

The principal finding is that the tested configurations could not be established as reliable single-cell reconstructions under a specific retrospective constraint: starting from an 8 µm bin-level Seurat object, without returning to Space Ranger or finer-resolution molecular inputs. Small regions can leave much of the original RNA unassigned, whereas expansion recovers more complete bin vectors without resolving their within-bin cellular origin. Downstream RCTD adds a complementary compositional description but does not remove this information limit. The inverse recovery–dominance association in Figure 3C highlights competing properties of the outputs; it does not identify a universally superior method. This is an evidence-limited operational conclusion, not a formal pass/fail classification based on validated cellular boundaries.

This intestinal context is important when interpreting the observed failure modes. Closely apposed nuclear profiles in compact epithelial or immune-rich fields can challenge separation, whereas elongated stromal and smooth-muscle profiles weaken a simple correspondence between nuclear geometry and the surrounding cell territory. Expansion may consequently cross anatomical interfaces while increasing RNA assignment. These are plausible anatomical explanations, not compartment-stratified findings of this benchmark: tissue compartments were not independently annotated, and boundary accuracy was not measured within them. The present case study therefore motivates anatomy-aware validation without establishing that intestinal complexity caused the differences between configurations.

The analysis used the deposited high-resolution H&E images, at approximately 1.3-1.4 micrometres per pixel. These inputs differ from finer-resolution whole-slide histology and from the training domains or preferred scales of some models. This comparison did not include systematic hyperparameter optimization: parameter configurations were fixed, a fluorescence-trained model was evaluated explicitly outside its intended imaging domain, and no annotation-based tuning was performed. Sparse output from a learned model should therefore motivate review of scale, domain and inference settings rather than a claim that the underlying algorithm is inferior.

Only two sections from one public study were examined. They are insufficient independent replicates for population-level inference. The Spearman correlation in Figure 3C is descriptive across 19 configurations, several of which are related parameter variants; it is not an inferential test across independent biological samples. Pairwise agreement in Figure 3A is conditional on bins fitted by both configurations, so high agreement can coexist with very different coverage. Dominance in Figures 3B–C is conditional on RCTD inclusion and can be affected by count depth, reference composition, transcriptional similarity and filtering. Because RCTD uses the same aggregated RNA whose recovery is being assessed, it provides complementary information rather than an independent validation of that aggregation. Without reference cellular boundaries or orthogonal identities, these quantities cannot establish segmentation or annotation accuracy. Manual inspection was unblinded and qualitative.

If only the deposited bin-level Seurat object is available, changing polygons cannot recover the unobserved within-bin allocation of RNA. Access to suitable original or finer-resolution inputs could permit a different upstream analysis, potentially including reprocessing with Space Ranger, but that workflow was not tested here and is not established as a remedy by these results. A broader validation would require independently annotated fields, prespecified nuclear and whole-cell tasks, orthogonal cell identities, additional tissue sections and controlled evaluation of image and molecular resolution. The present findings should not be generalized into a claim that no possible method can succeed, nor into a validated ranking of all segmentation algorithms.

## Conclusion

In this two-section Visium HD case study, no tested configuration could be established as a reliable single-cell reconstruction from the deposited 8 µm bin-level Seurat input. Higher RNA recovery did not imply better boundaries or greater RCTD dominance, and the latter did not establish correct cell identity. The practical conclusion is that segmentation-derived expression should not be treated as validated single-cell data on the basis of RNA recovery or RCTD support alone. This does not prove that all segmentation methods fail, that Visium HD cannot resolve cells, or that restarting from Space Ranger would necessarily resolve the limitations; upstream reprocessing was not tested. Under the available-input constraint, outputs should be reported as polygon-associated expression with explicit unassigned RNA and RCTD exclusions.

## Author contributions

Luca Massimino: conceptualisation, methodology, software, formal analysis, data curation, visualisation, writing – original draft, and writing – review and editing. Federica Ungaro: writing – review and editing.

## Ethics and use of existing data

This work was a secondary computational analysis of previously deposited expression data, spatial coordinates and associated H&E images from GSE302569 [1,2], together with the published scIBD reference [10]. No new participants were recruited, specimens collected or clinical interventions performed for this reanalysis. Zhang et al. report approval of the original study by the Medical Ethics Committee of Nanfang Hospital, Southern Medical University (NFEC-2024-495), and written informed consent [1]. The histology panels in this manuscript were generated from the deposited H&E inputs with newly computed analytical overlays, rather than copied from the published article’s figure panels.

## Generative AI assistance

OpenAI Codex was used to assist with code development and debugging, figure preparation, and manuscript drafting and language editing under human direction. The authors retain responsibility for verifying the analyses, references and interpretations and for the final manuscript. Histology panels were rendered from deposited H&E images and computed analytical outputs; no generative image synthesis was used.

## Data and code availability

The source study is indexed in the Gene Expression Omnibus under accession GSE302569 [2]; the scIBD reference is described by Nie et al. [10]. The analysis archive contains the 40 accepted segmentation masks, original bin-to-region mappings, gene-wise RNA-conservation audits, diagnostic metrics, all 38 successful native major-group RCTD fits, explicit zero-region and filter-exclusion records, and the data underlying Figures 1–3 and Supplementary Figures S1–S16. Versioned code snapshots, software and model-checkpoint manifests, source-coordinate and UMI provenance, checksums, and execution logs are retained. Deposited expression counts and author-defined reference labels were not altered. Analysis scripts, configurations, segmentation outputs, benchmark tables, figures and provenance manifests are publicly available at https://github.com/maximinio/zhang_segmentation_benchmark. Native RCTD result bundles and the deposited bin-level input object are available in the data-v1 release at https://github.com/maximinio/zhang_segmentation_benchmark/releases/tag/data-v1. File checksums and download links are provided in the repository manifests.

## Figure and supplementary table legends

Figure 1. Study workflow under a deposited bin-level input constraint. (A) Original H&E, counts and spatial coordinates are available from the deposited 8 µm Seurat object. (B) Twenty segmentation configurations are evaluated; whole integer bin count vectors are assigned to regions by bin centre, without molecule-level reassignment. The green boundaries illustrate H&E watershed D (+5 µm). (C) Native full-mode RCTD uses the scIBD RNAraw reference and author-defined major groups. Regions are coloured by the largest native coefficient; grey denotes regions excluded by native filters and black denotes exact tied maxima. The same preselected dense CD2 field and unchanged geometry are shown in all panels (scale bars, 100 µm). The example corresponds to Supplementary Figures S8 and S16 and illustrates the workflow, not a preferred method. Display calls are not calibrated probabilities or validation of cellular boundaries. No Space Ranger reprocessing or finer-resolution expression is introduced. Reference: Nie H, Lin P, Zhang Y, et al. Single-cell meta-analysis of inflammatory bowel disease with scIBD. Nature Computational Science. 2023;3:522–531. https://doi.org/10.1038/s43588-023-00464-9.

Figure 2. RNA assignment and region occupancy across all 20 configurations. (A) Percentage of the original section UMIs assigned by the common bin-centre rule. (B) Number of detected regions, in thousands. (C) Percentage of regions receiving no assigned RNA; this quantity is undefined for the zero-region microSAM LM AMG output. Blue circles and orange triangles denote CD1 and CD2; grey lines connect the two sections for each configuration. Nuclear, cell-target and expanded configurations are not biologically equivalent, and the fluorescence InstanSeg configuration is out of domain for H&E. RNA coverage and region counts do not measure boundary accuracy. A–D identify the marker-controlled watershed parameter configurations listed in Supplementary Table S1.

Figure 3. Cross-method concordance, ranked RCTD contributions and the RNA-recovery–dominance trade-off. (A) Pairwise agreement of dominant major-group calls on original 8 µm bins assigned to RCTD-fitted regions by both configurations. Each region’s largest native full-mode coefficient defines its display call; bin-level calls are compared only on the pairwise jointly fitted support. The heatmap is the arithmetic mean of the two slice-specific agreement matrices. Dendrograms use average linkage on 1 minus mean agreement. (B) Within each fitted polygon, the nine native coefficients are divided by their unmodified column sum for display and ranked from largest (Call 1) to smallest (Call 9), independently of cell identity. Rank contributions are averaged within each slice and then averaged equally across slices; bars are ordered by mean Call 1 contribution. This is a compositional concentration summary, not a count of accepted cell-type calls or a calibrated uncertainty probability. (C) One point per configuration pairs the arithmetic mean of slice-level assigned/original UMI fractions with the same mean Call 1 contribution shown in B. RNA recovery includes all segmented regions, whereas dominance is conditional on native RCTD inclusion. Colours identify methods. The descriptive Spearman correlation is −0.77 across 19 configurations; related method variants are not independent biological replicates and no inferential P value is reported. microSAM AMG produced no regions and is shown as undefined in B and omitted from A and C. Native coefficients were retained without clipping or refitting. No panel validates cellular boundaries or biological identity.

Supplementary Table S1. Complete diagnostic summary of the 20 configurations. RNA coverage is reported separately for CD1 and CD2 and pooled by summing original UMIs, not by averaging percentages. Counts of regions and area or occupancy diagnostics describe the output geometry. Targets differ across rows. Dashes denote undefined per-region statistics for configurations that detected no regions; these values are not measured zeros.

Supplementary Figures S1–S4. Segmentation outputs for tissue section CD1. Each row presents the complete section and the same high- and intermediate-cell-density fields across methods. Green outlines indicate all detected region boundaries. Scale bars: 1000 µm for complete sections and 100 µm fordetailed fields. RNA assignment percentages are calculated across the complete section, not the displayed fields. Figure S1 presents the five Cellpose configurations; Figure S2 presents both InstanSeg models, both microSAM LM configurations and PathoSAM; Figures S3 and S4 present the ten classical watershed configurations.

Supplementary Figures S5–S8. Segmentation outputs for tissue section CD2, using the same method order, field-selection criteria and scale-bar conventions as for CD1. Merged regions and long straight boundaries observed in some H&E watershed configurations are shown without selective exclusion. Greater RNA coverage does not establish accurate cellular boundaries.

Supplementary Figures S9–S12. Dominant RCTD major-group components for CD1, in the same method order and identical whole-section, dense-field and medium-density views as Supplementary Figures S1–S4. The accepted segmentation masks and original H&E pixels are unchanged. Each fitted region is coloured by the author-defined major_cluster group with the largest native spacexr 1.4.0 full-mode coefficient, using the scIBD RNAraw reference of Nie et al. [10]. This is a display call, not a calibrated posterior probability or a claim of pure-cell identity. Grey indicates regions excluded by native RCTD filters; exact tied maxima are shown separately. Fit counts and total regions are reported for each method. Scale bars are 1000 µm for whole sections and 100 µm for detailed fields. No confidence threshold, coefficient clipping or renormalization was applied; uncertainty is not quantified in these panels.

Supplementary Figures S13–S16. Dominant RCTD major-group components for CD2, corresponding to Supplementary Figures S5–S8 and using the same palette and display rule as CD1. The nine major groups are Myeloid, CD4T, CD8T, ILC, B_Plasma, Epithelial, Mesenchymal, Endothelial and Neural. microSAM LM AMG produced no regions in either section and therefore has no RCTD fit. All native fitted coefficients remain preserved in the analysis archive. These downstream compositional displays do not validate segmentation boundaries.

