## Supplementary Table S1 and Figures S1-S16 for "RNA assignment and segmentation geometry in Visium HD profiles of fibrostenotic Crohn’s disease"

#### Supplementary Table S1. Complete diagnostic summary and method configurations

| Method and target | Model and key parameters | RNA assigned (%) CD1 / CD2 / pooled | Regions CD1 / CD2 | Median area (µm²) CD1 / CD2 | Zero-RNA regions (%) CD1 / CD2 | Bin-corner discordance, UMI (%) CD1 / CD2 |
|---|---|---|---|---|---|---|
| Cellpose cyto3 cell candidates | cyto3; diameter 20 px; hematoxylin input. | 1.45 / 2.76 / 2.01 | 2,174 / 1,094 | 355.9 / 409.9 | 36.7 / 27.5 | 2.0 / 3.2 |
| Cellpose cpsam cell candidates | cpsam; diameter 20 px; RGB H&E. | 5.11 / 5.37 / 5.22 | 43,931 / 17,594 | 39.1 / 44.5 | 56.2 / 40.1 | 20.4 / 18.0 |
| Cellpose cpsam v2 cell candidates | cpsam_v2; diameter 20 px; RGB H&E. | 4.09 / 4.76 / 4.38 | 35,598 / 14,487 | 39.1 / 46.2 | 58.3 / 38.5 | 16.4 / 15.6 |
| Cellpose DINO small cell candidates | cpdino-vitb; diameter 20 px; RGB H&E. | 9.02 / 4.70 / 7.19 | 53,964 / 11,104 | 50.8 / 62.2 | 48.6 / 28.8 | 29.0 / 13.0 |
| Cellpose DINO cell candidates | cpdino; diameter 20 px; RGB H&E. | 6.04 / 3.04 / 4.77 | 42,902 / 9,247 | 41.1 / 51.6 | 52.5 / 33.3 | 23.6 / 9.9 |
| InstanSeg fluorescence cell candidates; fluorescence-trained model evaluated on H&E | fluorescence_nuclei_and_cells; RGB H&E; fluorescence-trained, out-of-domain. | 54.62 / 38.70 / 47.89 | 230,986 / 109,464 | 89.9 / 94.2 | 35.0 / 23.2 | 88.2 / 73.6 |
| InstanSeg H&E nuclei nuclear candidates | brightfield_nuclei; RGB H&E; calibrated pixel size. | 16.71 / 12.52 / 14.94 | 211,462 / 99,547 | 23.5 / 21.3 | 72.6 / 70.0 | 53.2 / 40.3 |
| microSAM LM AIS microscopy candidates; H&E domain mismatch | vit_b_lm; AIS; RGB H&E; microscopy model, domain mismatch. | 8.14 / 19.14 / 12.79 | 9,934 / 8,477 | 248.3 / 305.9 | 35.3 / 27.7 | 9.5 / 19.0 |
| microSAM LM AMG nuclear candidates | vit_b_lm; AMG; H&E converted to grayscale; fixed AMG thresholds. | 0.00 / 0.00 / 0.00 | 0 / 0 | — / — | — / — | 0.0 / 0.0 |
| PathoSAM H&E AIS nuclear candidates | vit_b_histopathology; AIS; RGB H&E; software-provided tiled inference. | 2.55 / 8.61 / 5.11 | 3,418 / 3,246 | 203.3 / 213.4 | 38.7 / 17.7 | 4.9 / 14.9 |
| scikit-image H&E nuclear candidates | scikit-image watershed; hematoxylin; σ 1 px; minimum object 15 px; seed separation 10 px. | 32.68 / 28.61 / 30.96 | 42,695 / 22,250 | 99.7 / 80.9 | 39.1 / 34.7 | 57.1 / 48.8 |
| OpenCV H&E nuclear candidates | OpenCV watershed; hematoxylin; σ 1 px; minimum object 15 px; seed separation 10 px. | 4.78 / 6.89 / 5.67 | 42,695 / 22,250 | 15.6 / 16.0 | 74.6 / 69.5 | 16.9 / 19.6 |
| H&E watershed A (nuclei) nuclear candidates | Hematoxylin; σ 0.7 µm; all holes filled; seed 2.5 µm (2 / 2 px); expansion 0 µm. | 30.98 / 28.92 / 30.11 | 291,533 / 140,149 | 21.5 / 19.6 | 73.4 / 71.1 | 65.4 / 53.3 |
| H&E watershed A (+5 µm) expanded nuclear candidates | Hematoxylin; σ 0.7 µm; all holes filled; seed 2.5 µm (2 / 2 px); expansion 5 µm. | 74.68 / 64.05 / 70.18 | 291,533 / 140,149 | 76.3 / 76.5 | 42.4 / 33.2 | 85.5 / 73.5 |

| Method and target | Model and key parameters | RNA assigned (%) CD1 / CD2 / pooled | Regions CD1 / CD2 | Median area (µm²) CD1 / CD2 | Zero-RNA regions (%) CD1 / CD2 | Bin-corner discordance, UMI (%) CD1 / CD2 |
|---|---|---|---|---|---|---|
| H&E watershed B (nuclei) nuclear candidates | Hematoxylin; σ 0.7 µm; all holes filled; seed 3.5 µm (3 / 3 px); expansion 0 µm. | 30.95 / 28.91 / 30.08 | 210,697 / 100,416 | 29.3 / 24.9 | 65.9 / 63.2 | 64.9 / 52.6 |
| H&E watershed B (+5 µm) expanded nuclear candidates | Hematoxylin; σ 0.7 µm; all holes filled; seed 3.5 µm (3 / 3 px); expansion 5 µm. | 74.63 / 64.03 / 70.14 | 210,697 / 100,416 | 109.5 / 112.0 | 33.9 / 23.6 | 83.8 / 71.2 |
| H&E watershed C (nuclei) nuclear candidates | Hematoxylin; σ 0.7 µm; holes <6 / <7 px filled; seed 2.5 µm (2 / 2 px); expansion 0 µm. | 29.10 / 26.78 / 28.12 | 308,393 / 149,465 | 21.5 / 19.6 | 72.6 / 70.1 | 68.4 / 56.7 |
| H&E watershed C (+5 µm) expanded nuclear candidates | Hematoxylin; σ 0.7 µm; holes <6 / <7 px filled; seed 2.5 µm (2 / 2 px); expansion 5 µm. | 74.57 / 63.77 / 70.00 | 308,393 / 149,465 | 74.3 / 74.7 | 42.2 / 33.5 | 89.3 / 77.9 |
| H&E watershed D (nuclei) nuclear candidates | Hematoxylin; σ 0.7 µm; holes <6 / <7 px filled; seed 3.5 µm (3 / 3 px); expansion 0 µm. | 29.07 / 26.76 / 28.09 | 222,174 / 106,576 | 29.3 / 26.7 | 64.8 / 61.9 | 68.2 / 56.3 |
| H&E watershed D (+5 µm) expanded nuclear candidates | Hematoxylin; σ 0.7 µm; holes <6 / <7 px filled; seed 3.5 µm (3 / 3 px); expansion 5 µm. | 74.52 / 63.75 / 69.96 | 222,174 / 106,576 | 107.5 / 110.2 | 33.4 / 23.7 | 88.0 / 76.0 |

Paired values are CD1 / CD2. Pooled RNA coverage is calculated from summed assigned UMIs divided by summed original section UMIs, not the mean of section percentages. Zero-RNA regions are retained. Dashes indicate undefined per-region statistics when no regions were detected. Bin-corner discordance quantifies sensitivity of bin-to-region assignment, not boundary accuracy.

For H&E watershed A–D: rgb2hed hematoxylin, Gaussian σ 0.7 µm, Otsu threshold within the common tissue mask, minimum foreground object size 6 / 7 pixels (approximately 12 µm² nominal), Euclidean-distance-transform local maxima, and scikit-image watershed on the negative distance map within foreground. A/B fill all internal holes; C/D fill only holes smaller than 6 / 7 pixels. Seed separations are nominal physical values with their effective integer-pixel values shown in each row. A nominal 4.5 µm setting is equivalent to 3.5 µm after rounding and is not an additional configuration. Expansion uses expand_labels, restricted to the common tissue mask. All outputs are computational candidates or territories, not independently validated complete-cell boundaries. RNA coverage alone does not establish biological segmentation accuracy.

#### Supplementary Figure S1

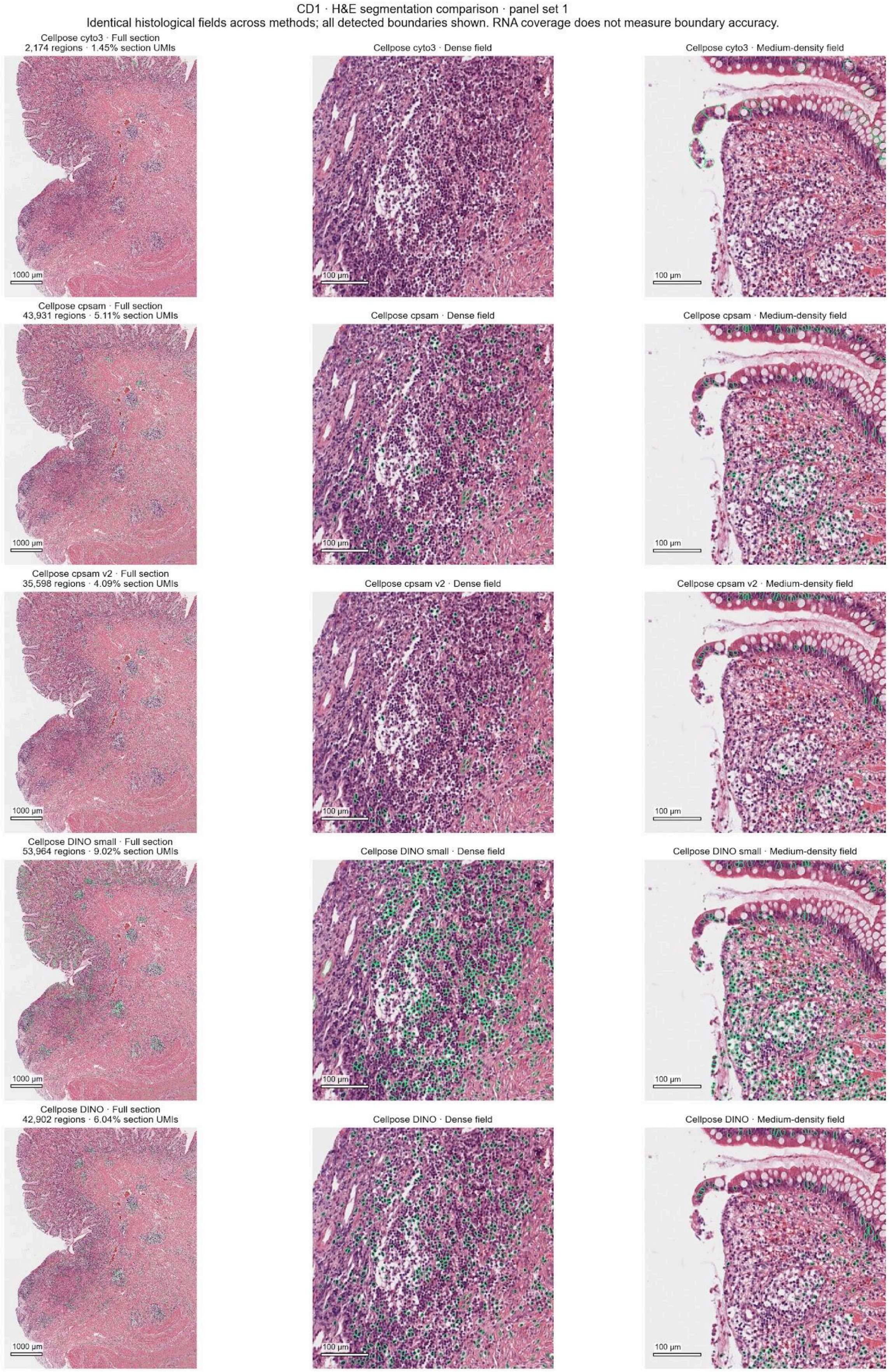

#### Supplementary Figure S2

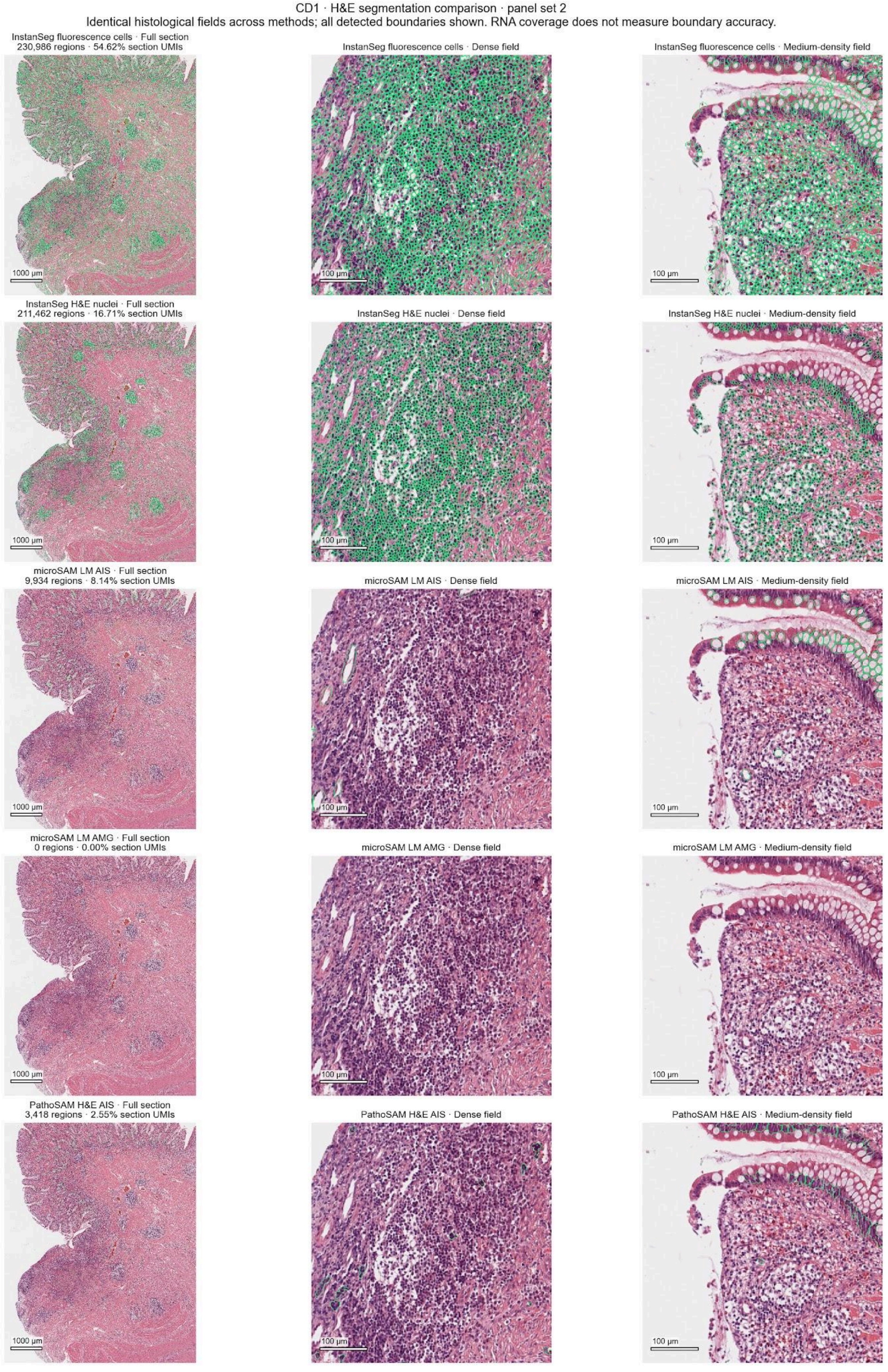

### Supplementary Figure S3

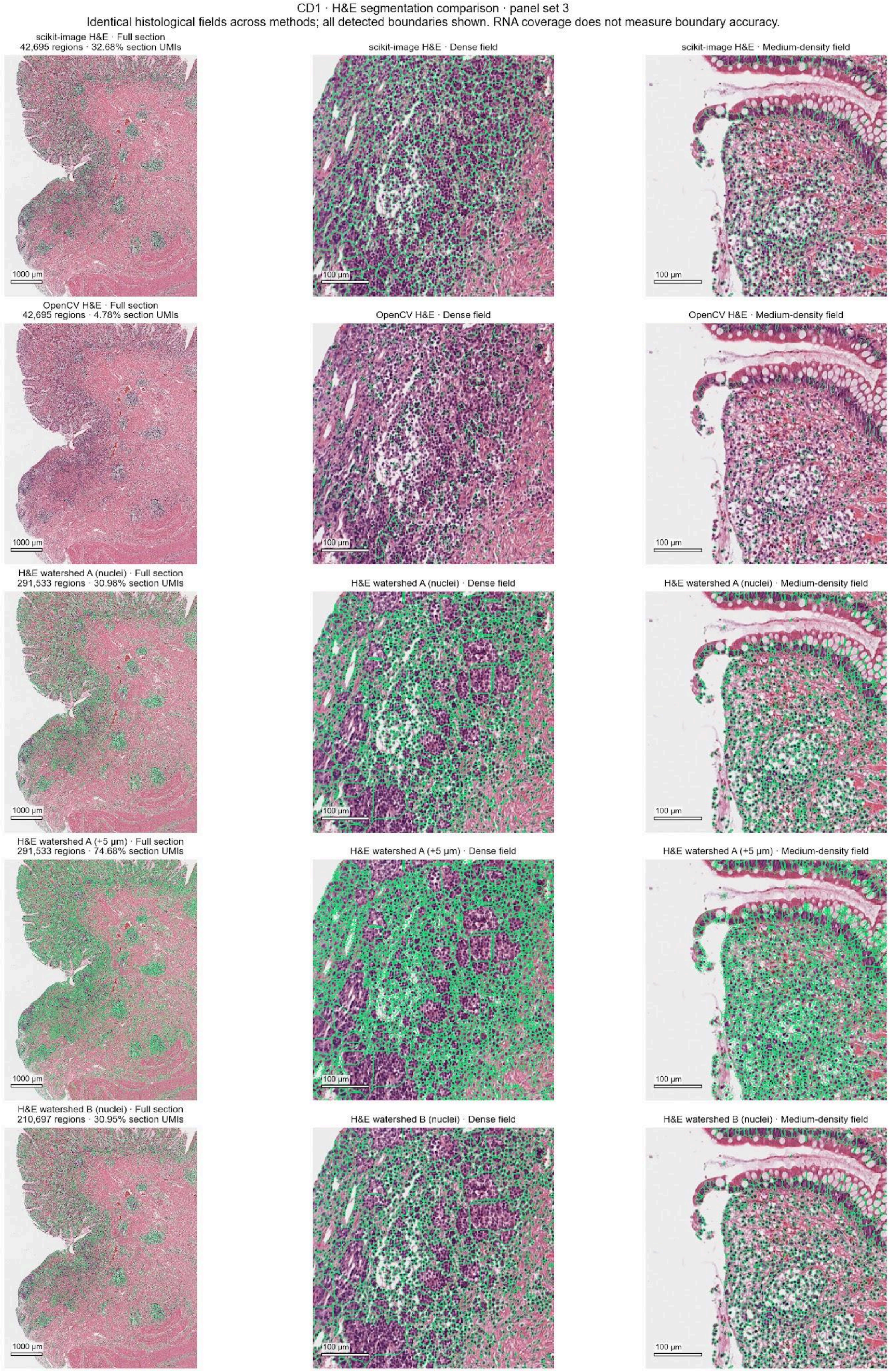

### Supplementary Figure S4

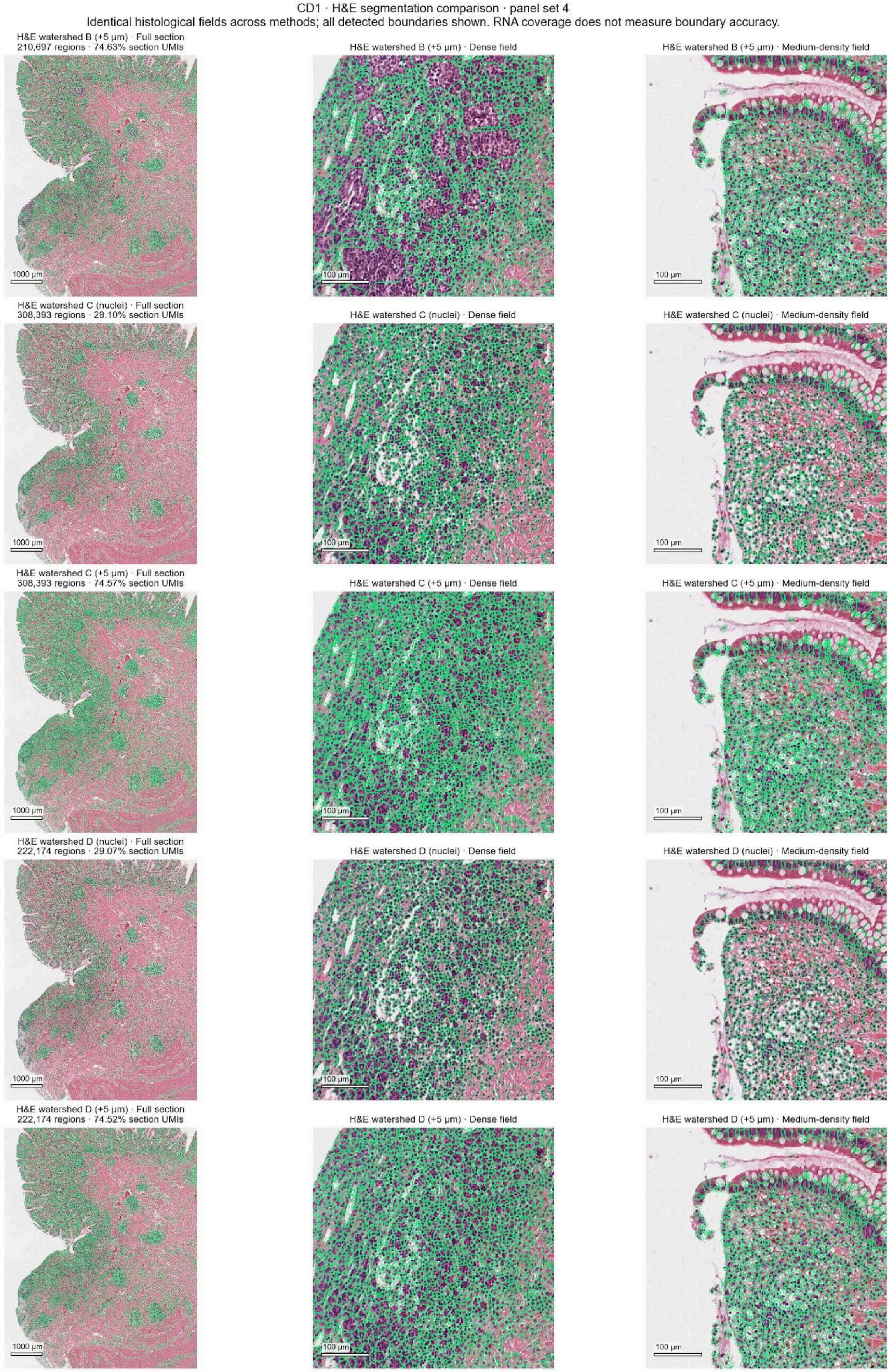

### Supplementary Figure S5

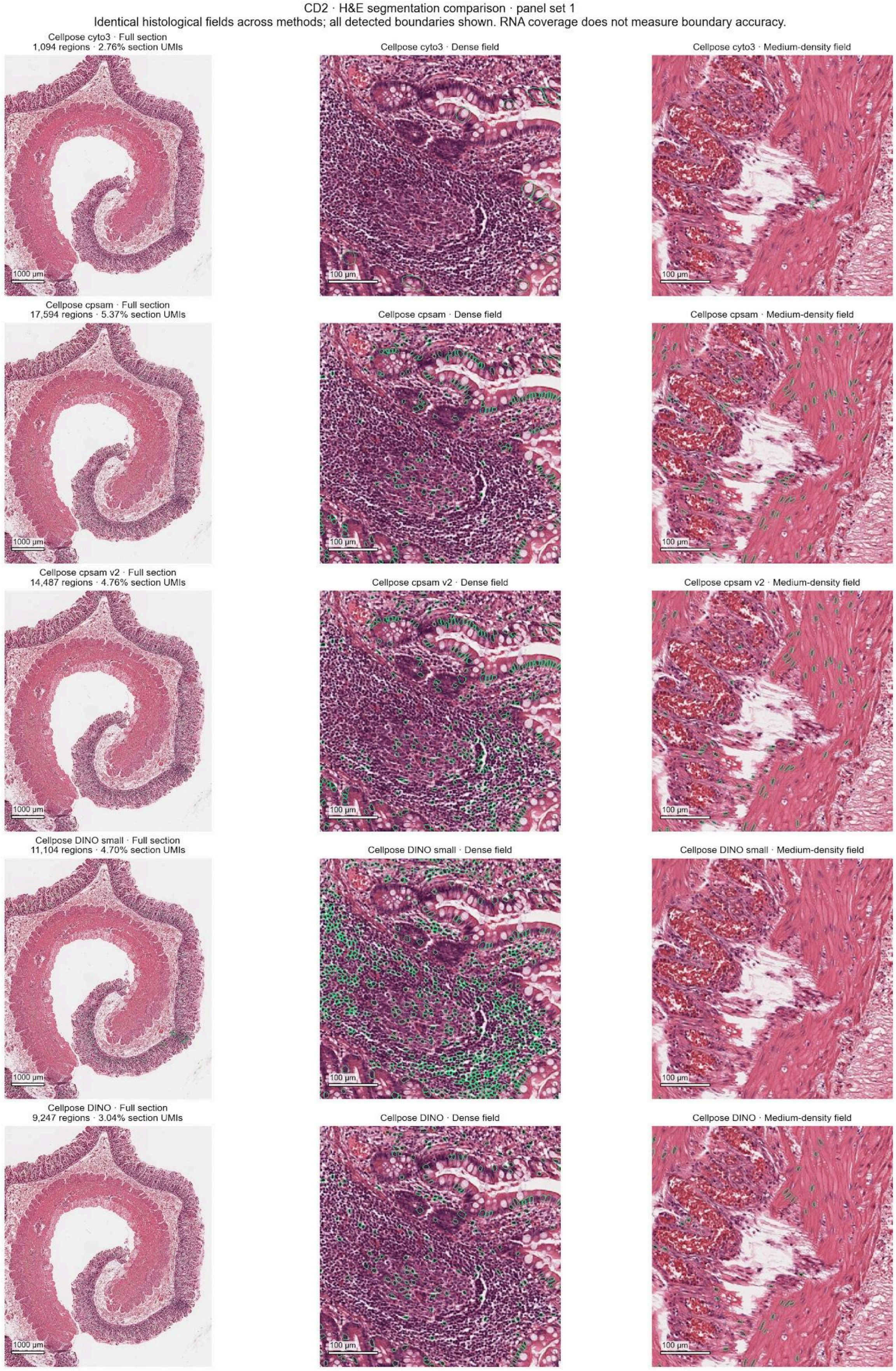

#### Supplementary Figure S6

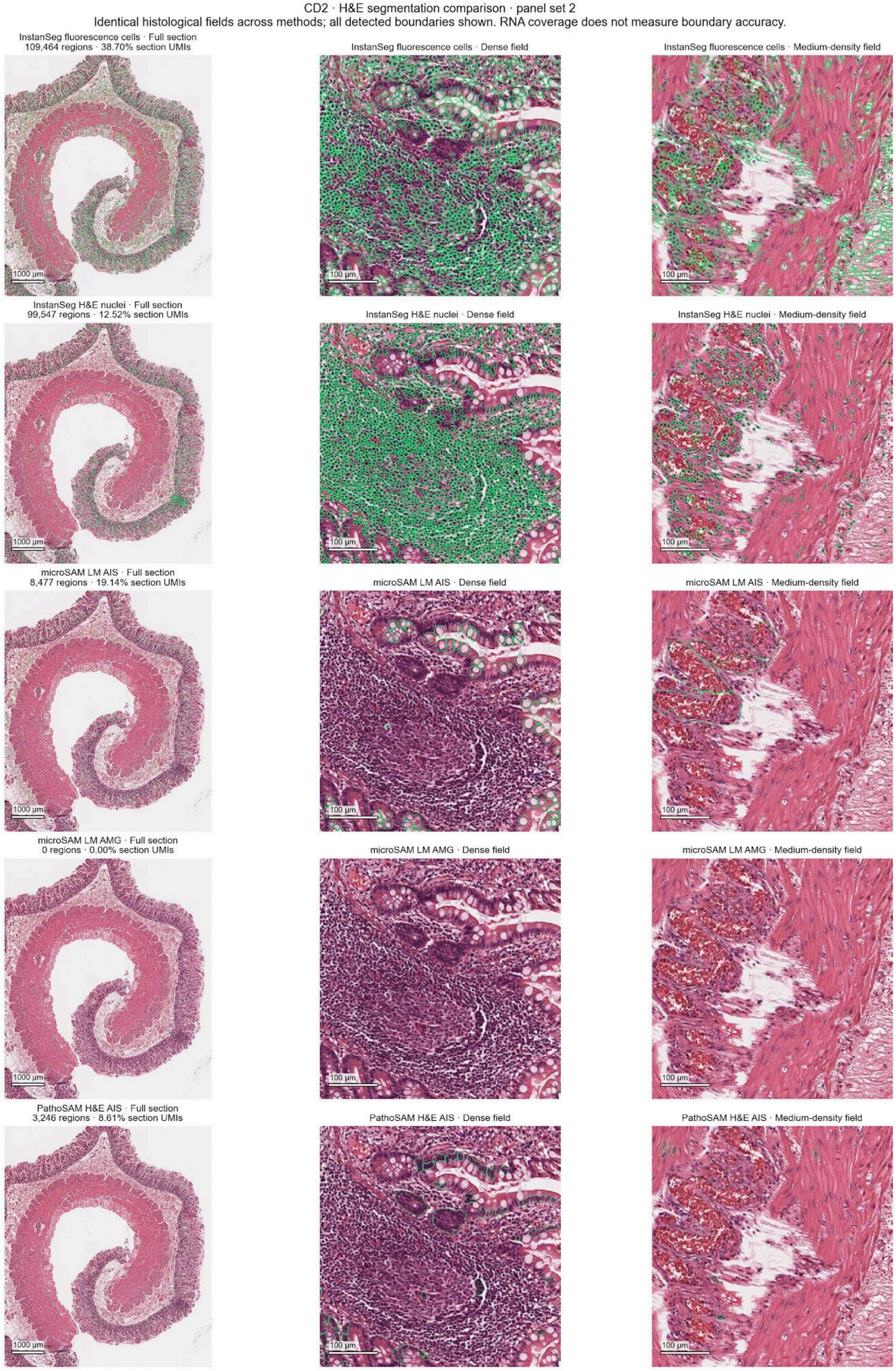

### Supplementary Figure S7

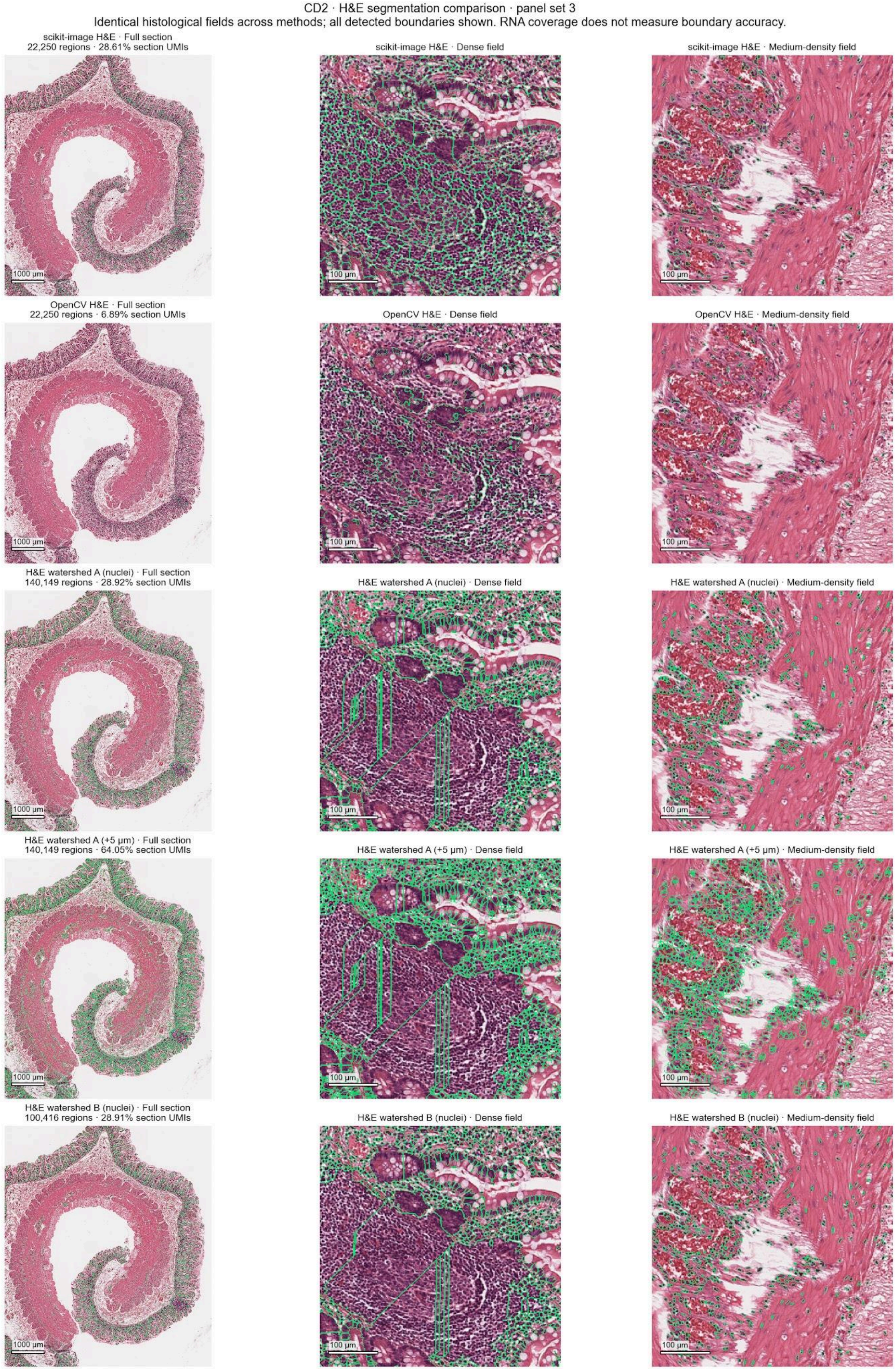

### Supplementary Figure S8

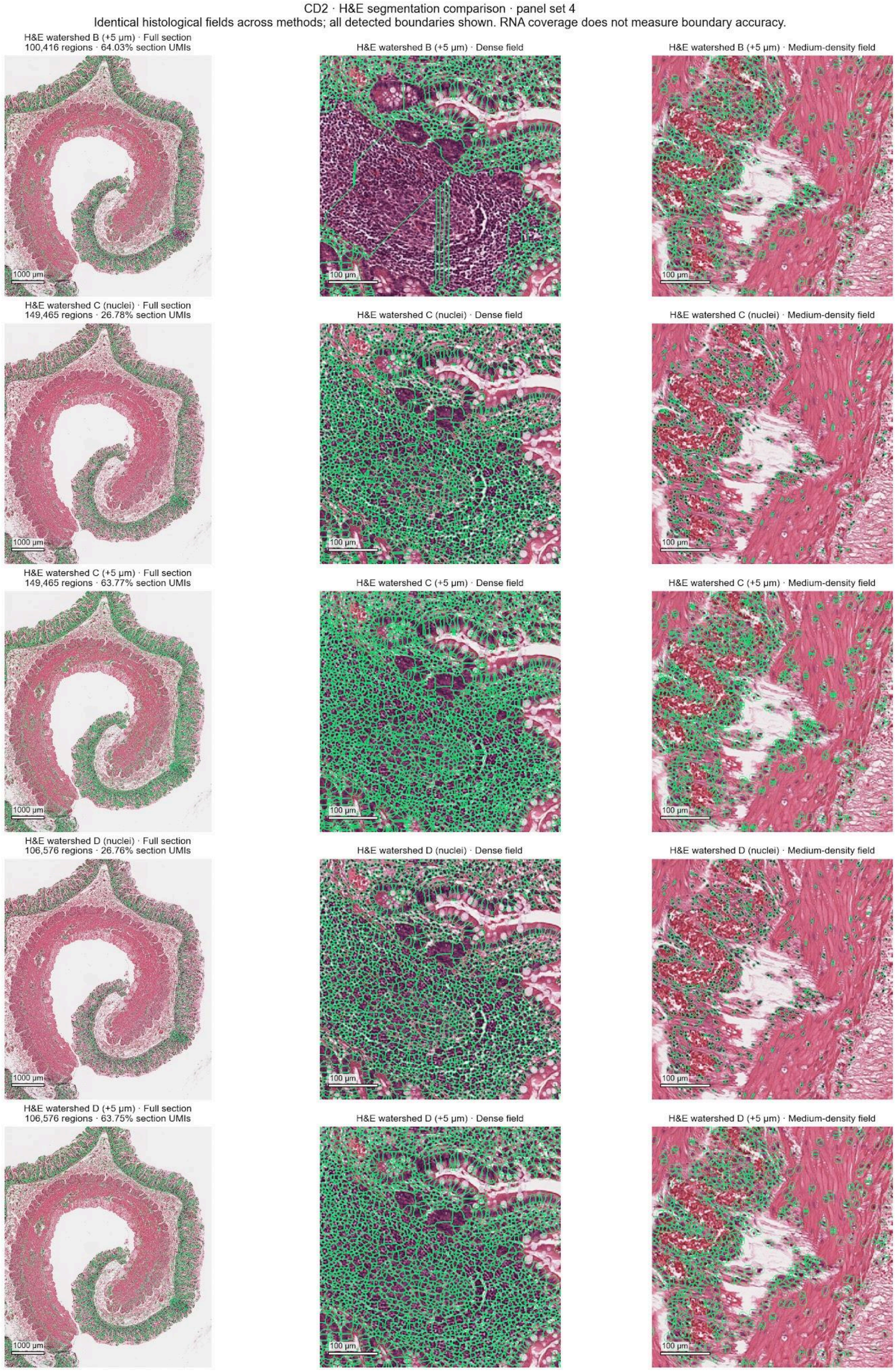

### Supplementary Figure S9

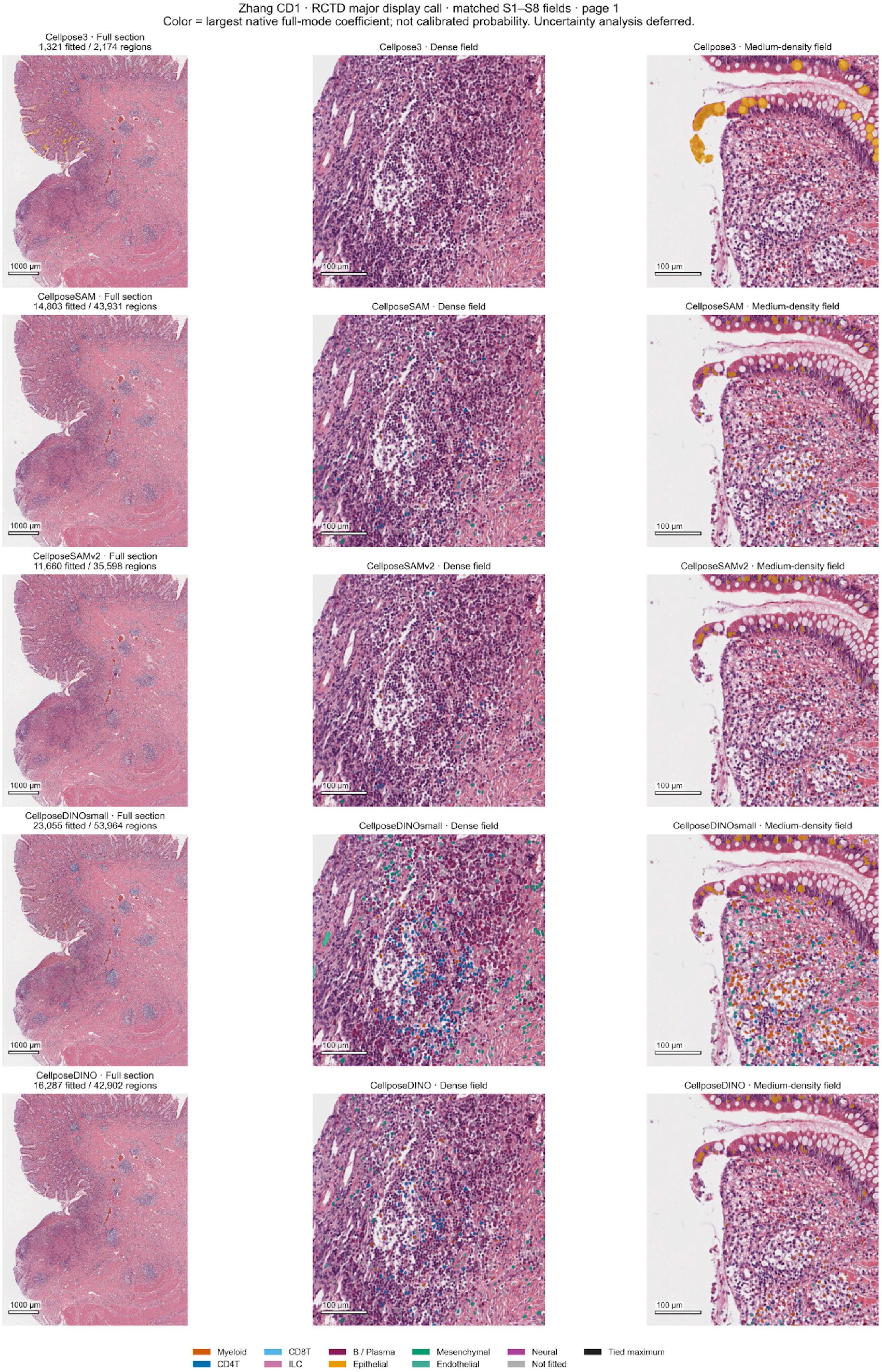

### Supplementary Figure S10

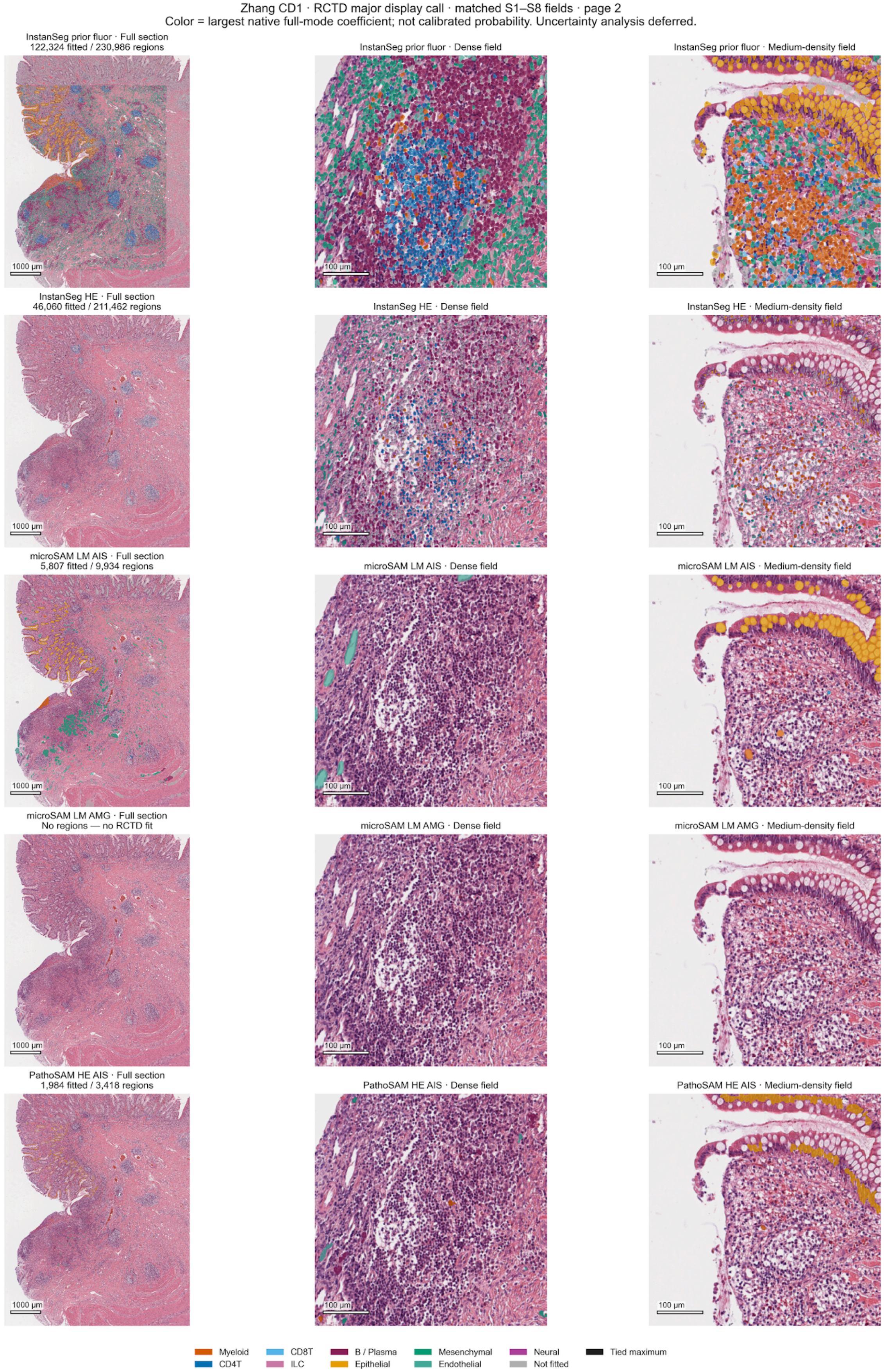

### Supplementary Figure S11

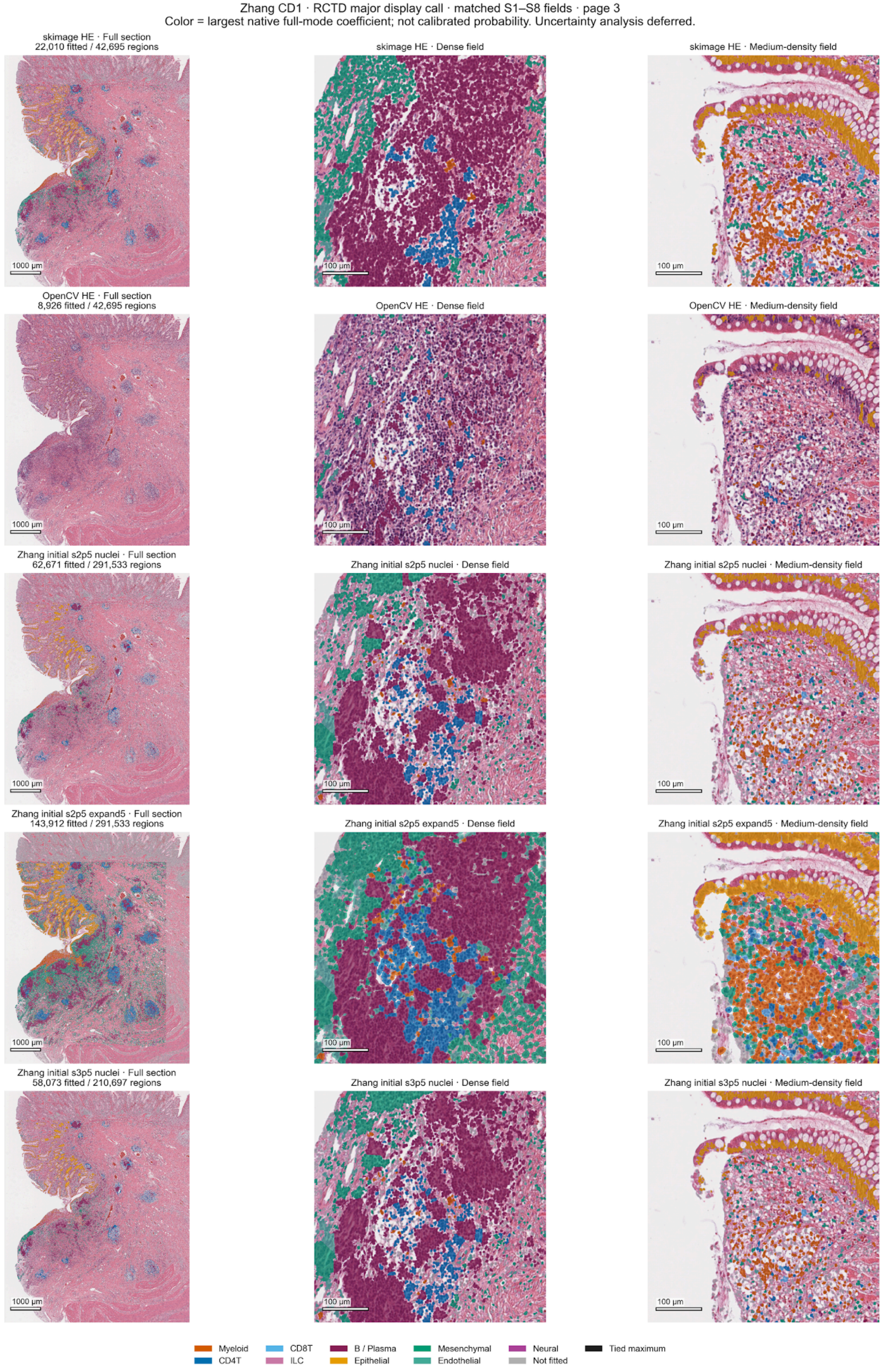

### Supplementary Figure S12

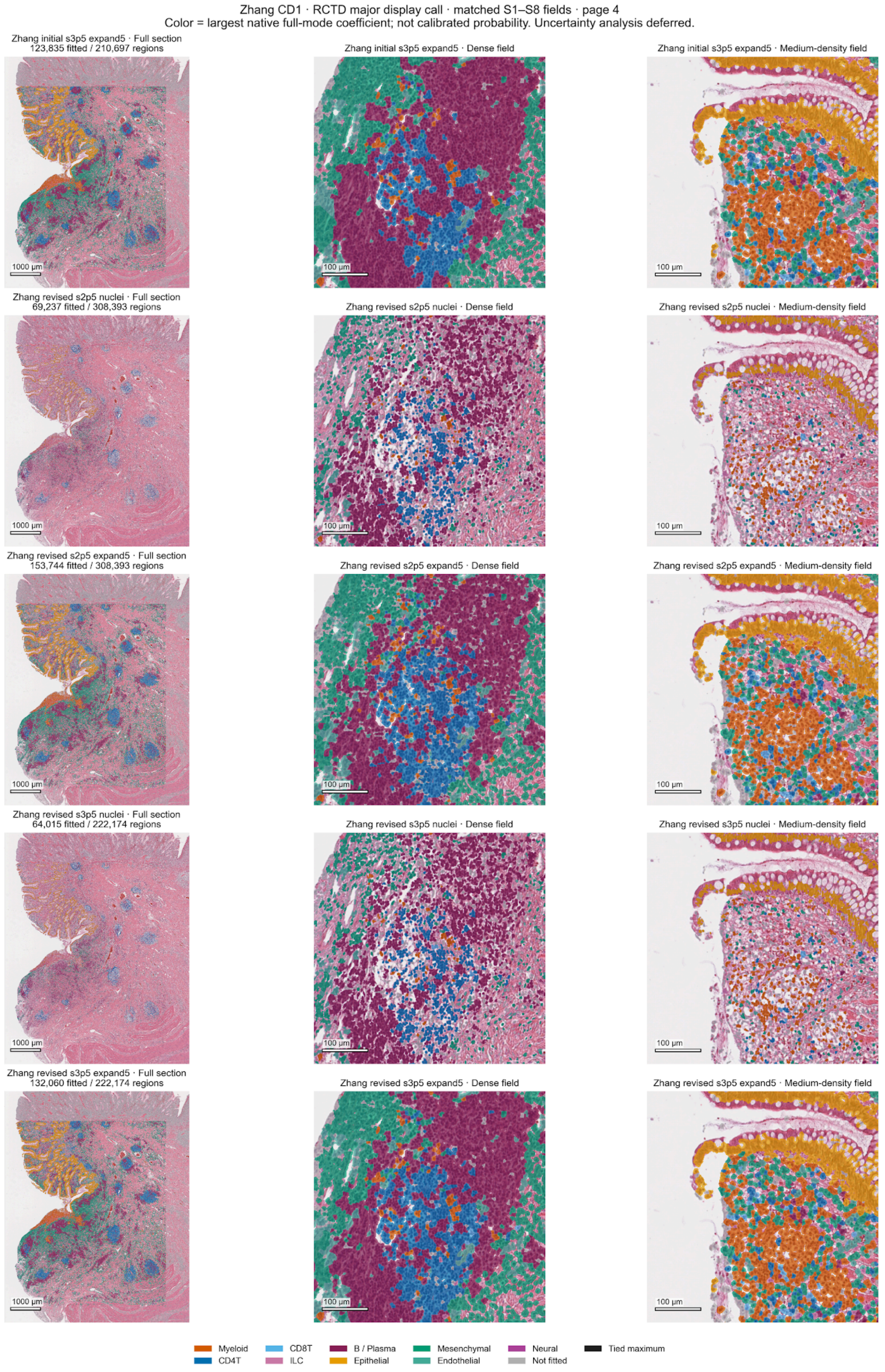

### Supplementary Figure S13

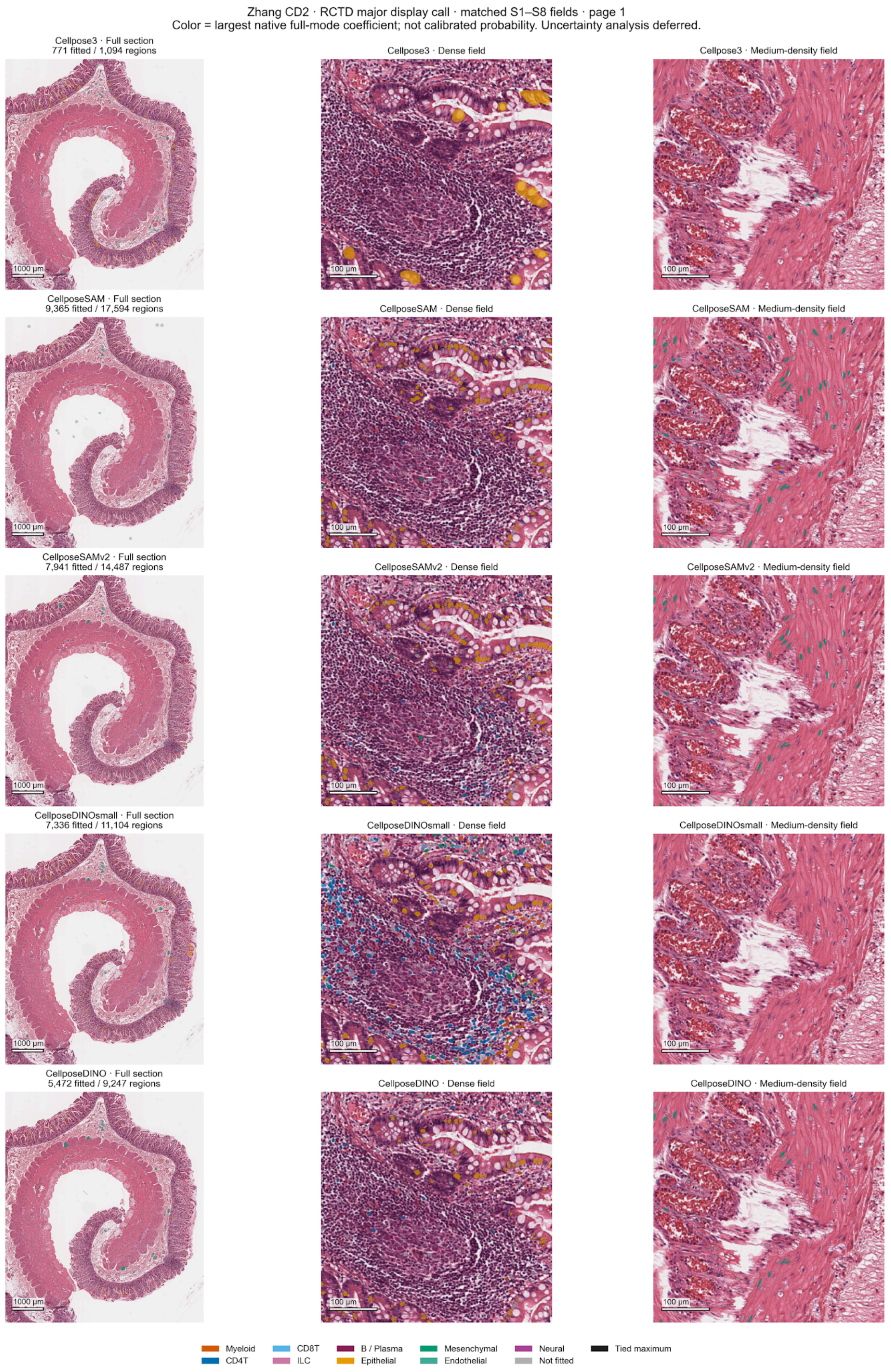

#### Supplementary Figure S14

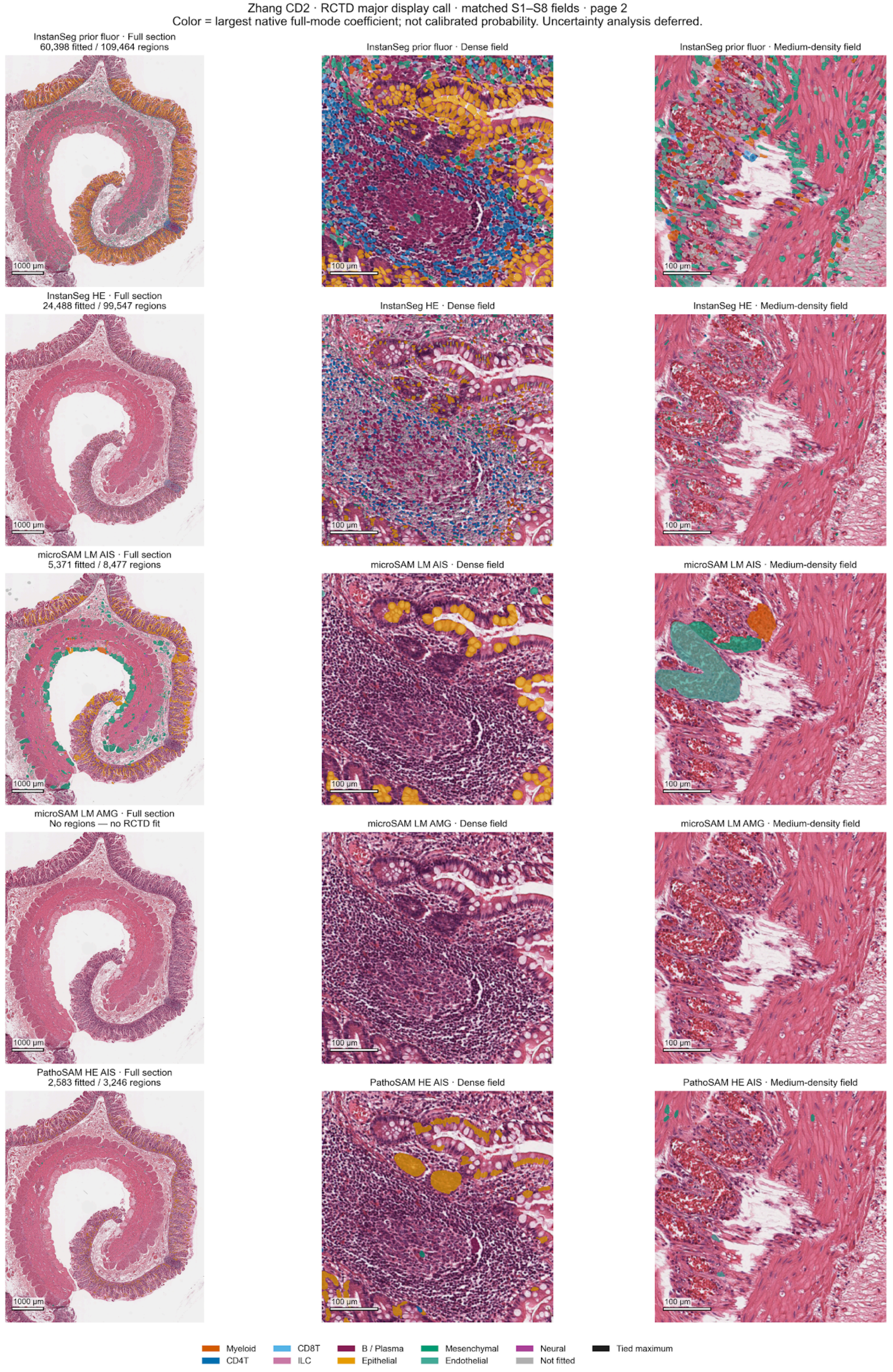

### Supplementary Figure S15

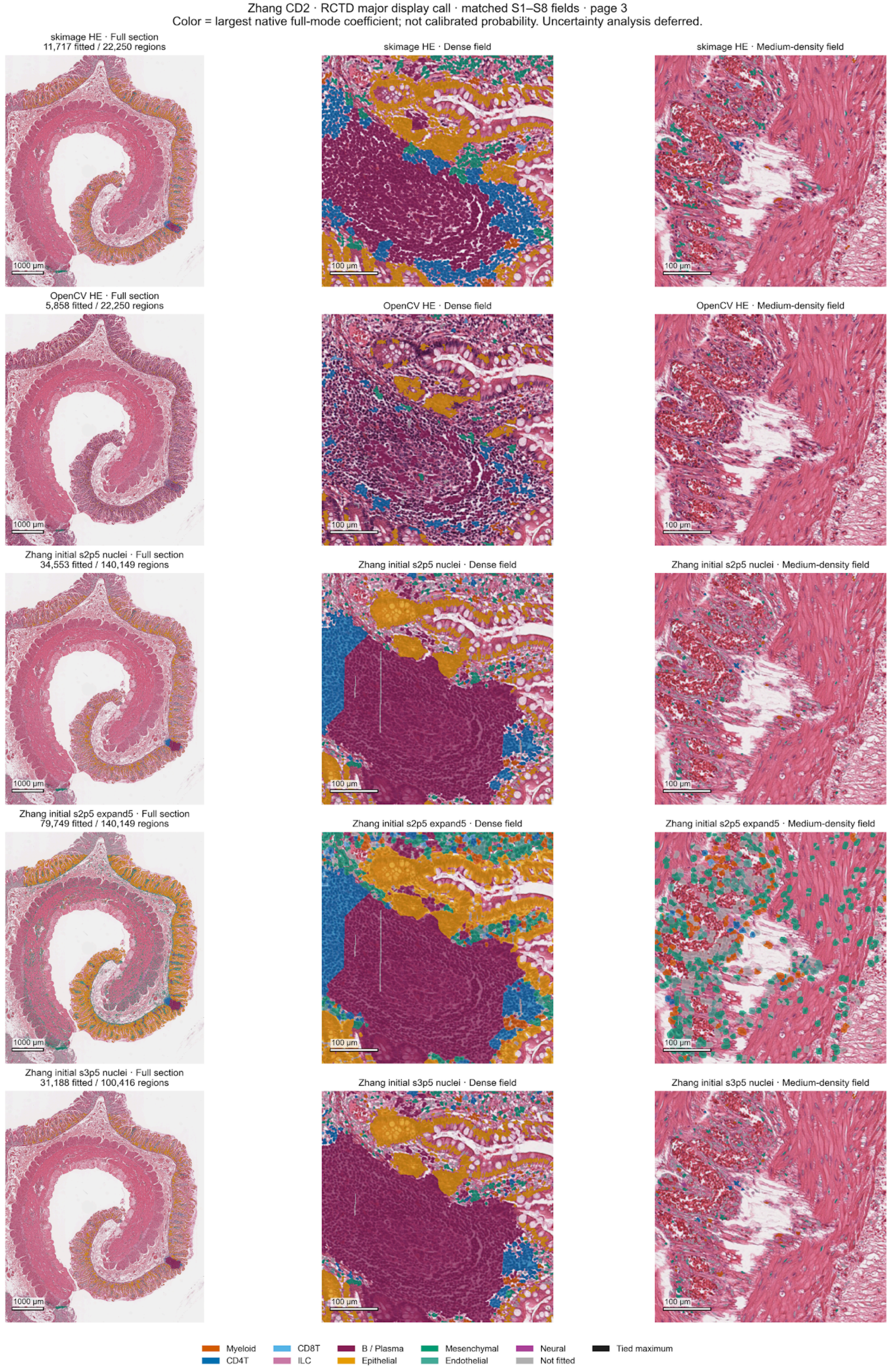

### Supplementary Figure S16

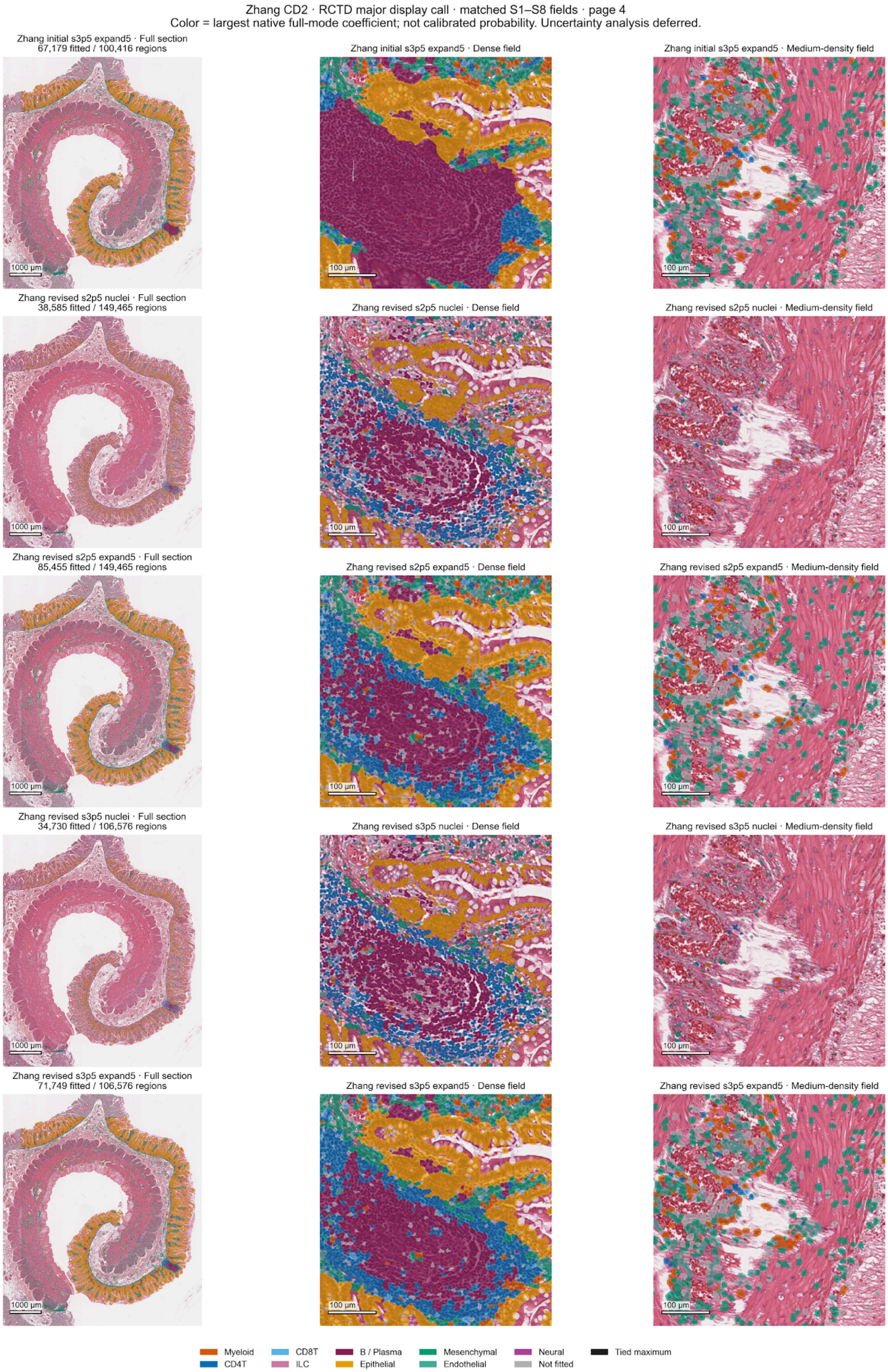
